# Correlative evidence for co-regulation of phosphorus and carbon exchanges with symbiotic fungus in the arbuscular mycorrhizal *Medicago truncatula*

**DOI:** 10.1101/618850

**Authors:** Jan Konečný, Hana Hršelová, Petra Bukovská, Martina Hujslová, Jan Jansa

**Affiliations:** Department of Experimental Plant Biology, Faculty of Science, Charles University, Viničná 5, Prague 2, 128 00, Czech Republic; Institute of Microbiology ASCR, Vídeňská 1083, Prague 4, 142 20, Czech Republic

**Keywords:** arbuscular mycorrhizae, barrel medic (*Medicago truncatula*), carbon, phosphorus, real-time PCR, shading, transcriptomics, transporter

## Abstract

In the research of arbuscular mycorrhizal (AM) symbiosis a considerable progress was made. But despite that, key questions still remain unanswered – for example it is well known that biotrophic fungus release phosphate (P) to- and recieves carbon (C) from the plant symbiont, but the particular genes, and their products, responsible for this exchange are still not fully understood. Here, we made a *de novo* quest for such genes involved in C transfer. Using physiological intervention of 90% shading and the correlation of expression levels of MtPT4, the AM-specific marker, and our candidate genes we demonstrate that several novel genes may be involved in AM symbiosis in *Medicago truncatula*. Also, we examined the expression of phosphate transporters (MtPT1-6) and we discuss the balance of “direct” and “mycorrhizal” P uptake pathways upon symbiotic fungus infection and C deprivation.

## Introduction

Research on the arbuscular mycorrhizal (AM) symbiosis is driven by continuous demand for understanding of this ancient symbiosis, and requirements to design more sustainable agricultural systems for the future. The arbuscular mycorrhizal symbiosis (AMS) is one of the oldest and the most widespread plant-microbe interaction on the Earth (Parniske, 2008) with a strong impact on the physiology and ecology of their plant hosts including many crop plants (Kim et al., 2017), as well as functioning and stability of entire ecosystems (van der Heijden et al., 1998, Wagg et al., 2014). In the times of rapidly changing environmental conditions, the influence of AM fungi on plant drought-, salt- or pathogen tolerance is growing in importance (Newsham et al., 1995; Bernardo et al., 2019 and references therein). Despite these features, it is still surprisingly little known about the economy of the symbiosis, i.e., the rates of exchanges of commodities between the symbiotic partners, and cellular changes in the host plants, particularly with respect to genes or proteins directly involved in metabolism and transfer of carbon (C) molecules towards the fungal partner (Garcia et al., 2016). When the AM fungus takes about 2-20% of plant C (Řezáčová et al., 2017), such knowledge seems important for understanding the physiology of AMS and C transfer in plants, as well as in understanding ecosystem processes particularly at times of high demand for increasing production and sustainability of agricultural systems (United Nations Sustainable Developmental Goals). This could be achieved either by direct application of AM fungi in agriculture, by utilization of indigenous AM fungal communities, or by getting specific „know-how“ of metabolic pathways used in AMS for improvement of crop yields and/or quality (Richardson et al., 2011; Hart et al., 2018).

Regarding phosphorus (P) exchanges between the symbiotic partners, the situation is rather simple. Plant can absorb P in the form of orthophosphate (mainly dihydrogen phosphate, H_2_PO_4_^−^) directly from the soil solution via phosphate transporters (PT) located on the rhizodermal cells (and their root hairs), with subsequent generation of P-depletion zones in the soil around the roots (i.e., “direct pathway”). To avoid the emerging P deficiency with plant increasingly exploiting P in the immediate vicinity of the roots, while relying solely on the direct P uptake pathway, a “mycorrhizal (indirect) pathway” is often established by a great majority of plant species (Smith et al., 2015). P is considered a major commodity transferred from the AM fungi towards the host plant, involving specific genes responsible for this pathway in periarbuscular membrane (PAM) of colonized root cells (arbuscocytes). In *Medicago truncatula*, MtPT4 was revealed as an AM-specific PT (Harrison et al., 2002; Smith et al., 2011; Jakobsen and Rosendahl, 1990), which is crucial for functional AMS in this plant species (Javot et al., 2007), making this gene one of the reference genes for established and functional AMS. Out of six PT described from the *M. truncatula genome*, MtPT1-MtPT6 (from here as PT1-PT6), it is the only one mycorrhiza-specific P transporter. Similarly, in other plant species we find a single or more AM-specific PT, too: LjPT3 (Maeda et al., 2006), OsPT11/13 (Paszkowski et al., 2002; Yang et al., 2012), StPT3/4/5 (Rausch et al., 2001; Nagy et al., 2005), LePT4/5 (Nagy et al., 2005) or HvPT8 (Glassop et al., 2005). During AMS, the expression of this AM-specific PT is highly upregulated, and consequently, the PT involved in the “direct pathway” tend to be downregulated (Liu et al., 2008). Yet, despite the extensive knowledge on P transfer from the AM fungus to the plant, much less is actually known about the mechanisms behind the reduced C transfer from the plant to the symbiotic fungus.

Biotrophic AM fungus acquires C solely from the host plant – this unique symbiotic behavior brings to mind an economic comparisons of C / P trading, however, the form as well as the rates of transferred C molecule are still a matter of debate (Raven et al., 2018). The most common hypothesis is a “sugar pathway”: export of sucrose (Suc) to periarbuscular space, where it is cleaved by acid invertases into glucose (Glc) and fructose (Fru), of which the Glc can be taken up by the fungal saccharide transporter (ST). This hypothesis is strongly supported by expression pattern and enzymatic activity of sucrose cleaving enzymes, mainly the cell wall-bound acid invertases (Salzer and Hager, 1991; Schaarschmidt and Hauser, 2008). Also, the substrate specificity of fungal ST (RiMST2, RiMST4 and RiMST5; Helber et al., 2011; Lahmidi et al., 2016), its expression in arbuscules and importance in formation of regular symbiotic structures such as arbuscules, supports the uptake of Glc and its relevance as one of the major forms involved in C transfer from the plant to the fungus. But the remaining Fru is generally not a substrate for those ST, so the fate of Fru is still unknown – leaving the question open about whether this form of C is taken back to the plant cells.

Another hypothesis is a transfer of fatty acid molecules towards the fungus. Higher production of fatty acids by the plants upon mycorrhization is logic, because newly arbusculated plant cell needs to synthetize new cell membranes – PAM is larger than the plasmatic membrane (Pumplin and Harrison, 2009) – but fatty acids may obviously also be a reduced C molecule transferred across periarbuscular space, as demonstrated recently (Rich et al., 2017a). Some key players in this “lipid pathway” were revealed, based on forward genetic screen of AM-defected mutants, and isotopolog profiling (Keymer et al., 2017). Notably, this cross-kingdom lipid transfer seems to play a major role for the physiology of the biotrophic fungus, since the dysfunction of lipid transfer prevents the completion of fungal life cycle by disrupted vesicle formation, reduced exploration of the root volume and incomplete development of extraradical mycelia and spores (Wang et al., 2012; Bravo et al., 2016; Bravo et al., 2017; Jiang et al., 2017; Keymer et al., 2017; Rich et al., 2017b; Brands et al., 2018), probably due to missing cytoplasmic fatty acid synthetase genes in *Rhizophagus genome* (Wewer et al., 2018).

Given the generally fragmentary knowledge about the molecular mechanisms of C transfer from the fungus to the plants, we describe here a *de novo* effort to identify genes potentially involved in symbiotic C transfer in the AMS, asking specific following questions:

1. which (novel) genes encoding ST – with particular focus on those ST showing mycorrhiza-specific transcriptional response – can play a role in C transfer from *Medicago truncatula* towards *Rhizophagus irregularis*, one of the model organisms in AMS research?
2. is there a transcriptional regulation when changing the availability of C for the plant, i.e., upon shading treatment, and shading-related alterations in P uptake pathways?

To achieve these goals, two pot experiments were conducted, in which the plants were grown for 35, 49, 63; 67 and 71 days, respectively, and where strong but short-term shading was applied on selection of the mycorrhizal and non-mycorrhizal control pots. Expression of genes was measured either by using Affymetrix microarrays and by specific quantitative real-time PCR assays. In addition to ST, we also measured the transcription of the different PT of the *M. truncatula* in both shoot and roots, and compared the expresion patters of genes involved in both mycorrhizal P and C fluxes.

## Materials and Methods

### Plant materials and growth conditions

We conducted two pot experiments (Exp 1 and Exp 2), using the same pot setup and glasshouse cultivation conditions, with an exception of the growing season and length of plant cultivation (for details, see below). Barrel medic (*Medicago truncatula* Jemalong 5; seeds originally provided by Gérard Duc from INRA, Dijon, France, and subcultured for several generations in the lab) seeds were surface-sterilized with 98% sulfuric acid for 20 min, rinsed thoroughly with distilled water and sown on wet filter-paper in room temperature. Two-days-old seedlings were planted in 2.5 L pots (3 or 4 seedlings per pot in Exp 1 or Exp 2, respectively) containing a mixture of autoclaved quartz sand (grain size < 3 mm), autoclaved zeolite MPZ 1-2.5 mm (Zeopol, http://www.zeolity.cz) and gamma-irradiated (> 25 kGy) LT soil (volumetric ratio 45:45:10; for soil characteristics see Řezáčová et al., 2016, for the characteristic of the planting mixture see Püschel et al., 2017) added with rhizobial suspension and with mycorrhizal or nonmycorrhizal inocula (for their description, please see below). The pots were placed in a glasshouse of the Institute of Microbiology (Czech Academy of Sciences, Prague, Czech Republic) at random positions with average temperature of 25/15°C during day/night. Light conditions were a combination of natural light and supplemental high pressure metal halide lamps (providing a minimum photosynthetic flux density of 150 μmol m^−2^ s^−1^ at plant level) to compensate for low natural light intensity and to extend the photoperiod to 14 h. Plants were watered daily with distilled water and fertilized weekly (starting 22 days post planting) with 65 mL of Long-Ashton solution with reduced phosphorus content (to 20 % of the original value, i.e., 0.26 mM P); Hewitt, 1966). In Exp 1, the plants grew from December to February and they were harvested as described below at 35, 49, 63 days post planting (dpp). In Exp 2, the plants grew from March to June and the plants were or were not placed under dark-green mesh-work canopy, allowing just 10% of the incoming light to pass through. Shading lasted for 4 or 7 days, after which period of time the plants were harvested (i.e., after 67 and 71 dpp; see Fig. 1).

**Figure 1:**
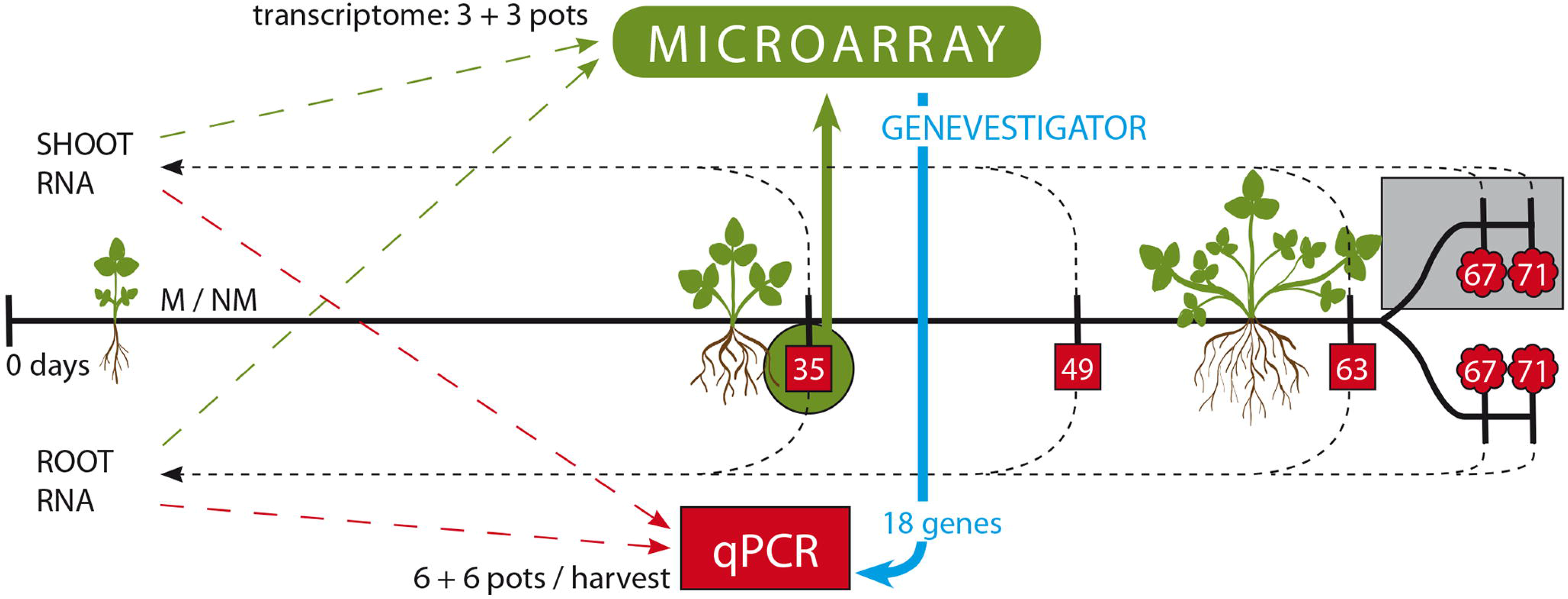
The scheme of both experiments. In Exp 1, the pots were harvested at 35, 49 or 63 days post planting (dpp), respectively. Whole shoot and whole root total RNA of 6 mycorrhizal (M+) and 6 non-mycorrhizal (NM) pots was isolated from plants collected at each harvest. From harvest at 35 dpp, the transcriptome was obtained (green, n = 3). Then, 18 genes (of those, 10 genes were newly selected via Genevestigator – blue) were subjected to qPCR evaluation (red), using all the harvests (red squares, n = 6). In Exp 2, 12 M+ and 12 NM pots were included, of which a half was shaded starting at 64 dpp, and all the pots were harvested for whole root and whole shoot total RNA at 67 or 71 dpp (i.e., 3 or 7 days after shading, if shaded; red clouds, n = 3).

### Rhizobial and mycorrhizal inocula

Rhizobial suspension of *M. truncatula*-compatible *Sinorhizobium meliloti* strain 10 (isolated from the LT soil; Püschel et al., 2017) was added to all pots before sowing (300 μL per planting hole/seedling, containing approximately 2 × 10^9^ cells). All the plants had well-developed nodules at the end of experiments (numbers and/or vitality not quantified).

Mycorrhizal inoculum was obtained from pot culture of an AM fungus *Rhizophagus irregularis* strain SYM5 (described in Gryndler et al. 2018), which was produced in the glasshouse for 8 months prior to the experiments described here, with leek (*Allium porrum*) as a host plant. Leek roots from the inoculum pots were harvested and cut mechanically into small pieces (< 1 cm) and then mixed back into the potting substrate from the inoculum production pots (which was the same as the potting substrate used in the experiments described here). This material (90 g) were mixed with 900 mL of the potting substrate mixture and such mycorrhizal (M+) inoculum was placed in the upper half of each of the M+ pots, so the pots in the M+ treatments contained cca. 5 % by volume of the mycorrhizal inoculum per pot. Non-mycorrhizal (NM) treatments were prepared in the same way, but the NM inoculum (also called mock inoculum) devoid of the AM fungus but produced for the same period of time under the same conditions as the M+ inoculum, was added to the NM pots instead of the M+ inoculum. Absence of mycorrhizal fungus in the mock inoculum pots was checked microscopically and also by qPCR (using the mt5 marker set described by Couillerot et al., 2012, data not shown) before application to the relevant pots.

### Plant harvest and RNA isolation

The plants from both experiments were harvested the same way, but the time of harvest was between 11 and 12 a.m. and between 1 and 2 p.m. in Exp 1 and Exp 2, respectively. All plants in one pot were processed as a single unit. First, the shoots were cut at the hypocotyl-root boundary and immediately frozen in liquid nitrogen. Thereafter, the roots were quickly recovered from the substrate, cleaned by shaking and under running cold tap-water, then the whole root system was blotted against paper towel and immediately frozen in liquid nitrogen. Plant samples were subsequently ground in mortar (while kept frozen at all times) and then subjected to RNA isolation or powderized samples further stored frozen at −80°C.

Frozen and pulvenized plant tissue – be it shoots or roots (cca 3 g) – was mixed with 4 mL of phenol and 4 mL of extraction buffer (1.95 g of 100mM Tris, 0.0424 g of 100mM LiCl, 0.372 g of 10mM EDTA and 1 g of 1% SDS per 100 mL of DEPC-treated water (0.1% DEPC), adjusted to pH = 6), pre-heated to 80°C, eight zirconium (ZrO_2_) beads were then added and the mixture was vigorously vortexed. This was followed by 20 s incubation at 80°C in a water bath, followed by 40 s vortexing, and mixing with additional 4 mL of chloroform. The mixture was then centrifuged (20 min, 0°C, 5000 g) and supernatant was carefully recovered. Addition of chloroform and centrifugation (purification steps) were repeated four-times. Pure supernatant (2 mL) was mixed with 0.68 mL of 8M LiCl. Samples were precipitated on ice overnight and then centrifuged (20 min, 0°C, 20000 g). The pellet was resuspended in 0.5 mL of 75% EtOH (ice-cold). The samples were centrifuged again, the pellets then diluted in 250 μL of RNase-free water, 500 μL of 96% ethanol and 25 μL of 3M sodium acetate. Samples were then precipitated on ice for 1 hour, then they were centrifuged and the pellets were diluted in 100 μL of RNase-free water. Resulting total RNA was frozen in liquid nitrogen and stored at −80°C.

### DNase treatment and reverse transcription

To remove DNA from the total RNA samples, DNA-free™ DNA Removal Kit (Ambion, USA) was used following the instructions of manufacturer. To synthetize complementary DNA strand, Transcriptor High Fidelity cDNA Synthesis Kit (Roche, Switzerland) was used following the instructions of manufacturer, but using both primer types served at once: 2 μL of Random Hexamer Primer (600 pmol/μL) + 1 μL Anchored-oligo[dT]_18_ Primer (50 pmol/μL). A maximum of 4 μg total RNA was used as template. Resulting complementary single-strand DNA (cDNA) was stored at −20°C.

### Gene expression analyses on Affymetrix

Total RNA of roots and shoots of three M+ and three NM pots, grown for 35 days, was subjected to transcriptome-wide gene expression analysis using commercial Affymetrix microarrays for *M. truncatula*. The analyses were carried out in one of the European certified Affymetrix core lab (http://core.img.cas.cz) using GeneChip™ Medicago Genome Array (Applied Biosystems™, USA). The RNA quality was checked prior to the analyses by RNA-gel electrophoresis and Agilent Bioanalyzer 2100 (data not shown) and raw data were then curated and uploaded to Genevestigator^®^ 4-36-0 (Nebion, Switzerland) database (Hruz et al., 2008). Those experimental data were also deposited in NCBI’s Gene Expression Omnibus (Edgar et al., 2002) and are accessible through GEO Series accession number GSE126833 (https://www.ncbi.nlm.nih.gov/geo/query/acc.cgi?acc=GSE126833).

Application “Samples” within the Genevestigator platform was used to search through by key-words (“sugar”, “transmembrane”, “transporter”, “saccharide”, “monosaccharide”, “sucrose”, “sucrose synthase”, “symporter”, “antiporter”, “alpha beta”, “glucosidase”, “glucose”, “glycan”, “fructose”, “fructan”, “trehalose”, “sorbitol”, “mannitol”, “mannose”, “mannan”, “ascorbate”, “cellulose”, “sweet”, “invertase”, “sucrase”) and also to seek through the published lists of ST (JCVI Medicago truncatula annotation database – Tang et al., 2014; Doidy, 2012 – for details see Table S1). Ten genes returning a clearly visible changes in their expression levels in roots or in shoots between M+ and NM plants were manually selected and briefly analyzed for their relevance in saccharide transport or metabolism. One reference gene (TIF2Fα) was identified using “RefGenes” application (Hruz et al., 2011; Table S1).

### Primer design

For analysis of gene expression by qPCR of the phosphate transporters PT1 through PT6 and previously used TEF1α reference gene, the published primer sequences were used (Grunwald et al., 2009, Baier et al., 2010; see Table S1). For 10 genes selected from the Microarray analyses, and the novel reference gene TIF2Fα, the qPCR primers were designed using AlleleID 6.0 (Premier Biosoft, USA; see table S1). Target sequences of primers were 100% identical to the target sequences of corresponding Probe Sets of GeneChip^®^ Medicago Genome Array. The primers were subsequently synthetized and HPLC purified by Generi Biotech (Hradec Králové, Czechia).

### Quantitative PCR

Quantitative PCR reactions included 13.2 μL of water, 0.4 μL of forward primer, 0.4 μL of reverse primer, 2 μL of cDNA template and 4 μL of 5× EvaGreen^®^ mastermix (Solis BioDyne, Estonia). Reaction conditions were as follows: initial denaturation at 95°C for 15 min, then 50 cycles of 95°C for 10s, annealing at 52-60°C (for details see Table S1) for 1 min and amplification at 72°C for 10 s. The analyses were carried out in StepOnePlus™ Real-Time PCR System (Applied Biosystems, USA).

Respective amplicons were purified by QIAquick PCR Purification Kit (Qiagen, Germany) and their lenghts were verified by DNA gel electrophoresis (data not shown). From the molecular weight of the fragments and DNA concentrations measured by Quant-iT™ PicoGreen™ dsDNA Assay Kit (Invitrogen™, USA), the concentrations of individual amplicons (in copies per microliter) were calculated. The amplicons and the information on their concentrations were used for calibration of the different qPCR assays on the same plate (using serially diluted amplicon preparations).

### Calculations and statistics

For statistical analyses of the results, we used programming language R 3.0.0 in RStudio 0.99.902 (RStudio, international) environment. Command “t-test” was used for Welch Two Sample t-test comparison, “lm” and “anova” for linear models and analyses of dispersion, and graphic commands “lineplot.CI” and “xyplot” for imaging the data. Due to inconstant expression levels of the reference genes, the results are not normalized against the reference genes, but they are normalized against the amount of total RNA subjected to reverse transcription of each corresponding sample.

### Analyses of phosphorus content

In Exp 1, the plant biomass was oven-dried at 65°C to constant weight. Dried samples of root and shoot biomass was ground to powder using a ball mill (MM200, Retsch, Haan, Germany) and of that, 0.08 – 1 g was used for further analysis. Milled samples was incinerated in a muffle furnace at 550°C for 12 h and the resulting ash was briefly heated to 250°C on a hot plate with 1 mL of 69% HNO_3_. The materials was then transferred to volumetric flasks through a filter paper and brought up to 50 mL with ultrapure (18 MΩ) water. Phosphorus concentration in the extracts was then measured by colorimetry at 610 nm using a Pharmacia LKB Ultrospec III spectrophotometer by the malachite green method (Ohno and Zibilske, 1991). Total P content was calculated from shoot- and root dry weight data and the concentrations of P in shoot and root biomass, respectively.

## Results

### Mycorrhizal plants have higher P content, but do not differ in biomass from the NM plants

In Exp 1, we have performed several paralel harvests for biomass, P and root colonization by the AM fungi assessments, in addition to the plants harvested for RNA analyses (as described above). The overall appearence of NM and M+ plants was consistent, which is confirmed by dry total biomass, dry root biomass, dry shoot biomass and root to shoot ratio (Fig. S1). It is important to mention that M+ treatment resulted in fully developed AMS, as also observed in the expression of AM-specific PT4 transporter (see Results bellow), by occurence of AM fungal structures including arbuscules in the M+ roots (Fig. S3). A strong difference in P concentration between the M+ and NM treatments was recorded. At 35 dpp, the M+ plants have nearly double the concentration of P in their tissues (total, root and shoot P concentration) and regarding the harvest at 64 dpp, the total content of P in the M+ plants was more than double of that in the NM plants (Fig. S2).

### Mycorrhizal phosphate-uptake pathway is active in M+ plants and reacts to shading

The AM-specific phosphate transporter (PT4) was expressed specifically in the M+ roots of *Medicago truncatula* (Fig. 2), while other (direct phosphate-uptake pathway) transporters (PT1, PT2 and PT3) were downregulated by the presence of AM fungus. In Exp 1, this pattern was consistent in all time points, with exception of 63 dpp for the PT2. In M+ plants, the number of PT4-transcripts was about four-times higher than those of the other PTs. The expression of PT4 was thus strongly dominating the PT-transcript-pool in M+ roots with relative proportion of 60-90 % of the four PT (direct + mycorrhizal pathways). In Exp 2, short-term light deprivation only had a little effect on expression values of the PTs involved in the direct P-uptake pathway, of which only PT2 was significantly upregulated in shaded NM plants, but downregulated in other time points (Fig. 2).

**Figure 2:**
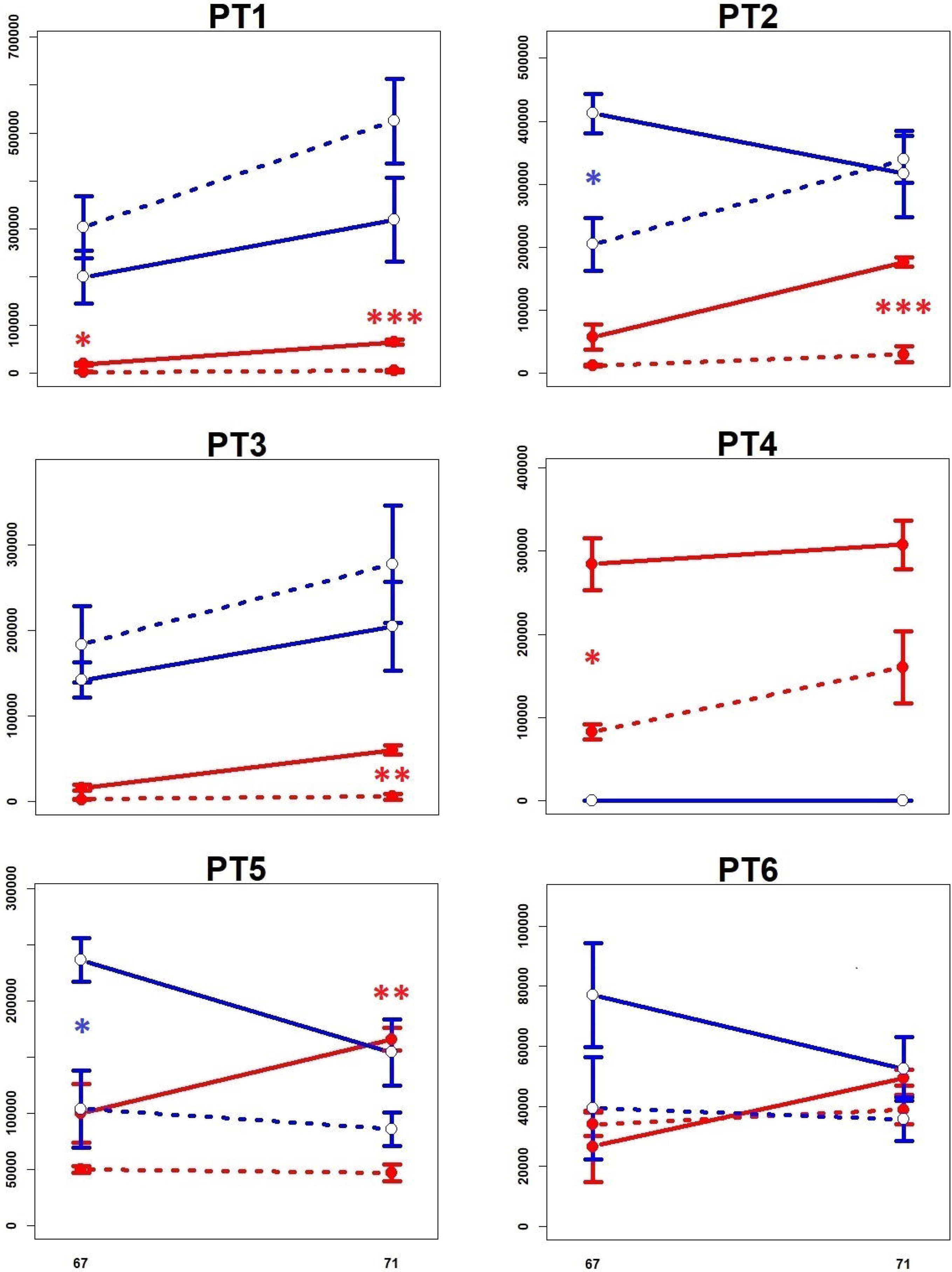
The expression of genes for high-affinity inorganic phosphate transporters (PT1 – PT6) obtained by qPCR from roots of *M. truncatula* in Exp 2. X-axis: days post planting, Y-axis: number of corresponding-gene copies per 1 µg RNA subjected to reverse transcription. Red: mycorrhizal, Blue: non-mycorrhizal, Full line: full light (100%), dashed-line: shaded plants (10% of light). Error bars show standard deviation, n = 3. For further details see Materials and methods and Table S1. Signif. codes (p-values) for t-test between non-/shaded: 0 < *** < 0.001 ≤ ** < 0.01 ≤ * < 0.05 ≤ non-significant.

On the contrary, the mycorrhizal pathway (PT4) was strongly downregulated upon 90% shading in Exp 2 (Fig. 2). Regarding number of copies measured, the expression of PT4 was about three-times lower in shaded M+ plant roots than in their unshaded M+ counterparts, but still a dominant PT-transcript of all six homologues.

PT5 and PT6 played a dominant role in shoots (Fig. S4), while the proportion of their transcripts to all six PT stably remained on the same high level (88-99 %, 95 % on average) in shoots, regardless of the environmental conditions. It is important to mention that PT5 and PT6 were significantly downregulated in M+ shoots from Exp 1 as compared to NM shoots. In roots, the ratio of PT5- + PT6-transcripts to all PT-transcripts was just around 22 % on average, regardless of the other variables (harvest time or shading).

### Some genes encoding saccharide transporters exhibit altered expression status in M+ roots

Using a manual-search in the whole-transcriptome data, 10 genes were selected for further analyses, which exhibited altered level of transcription upon mycorrhization. Of the available list of *Medicago truncatula‘s* saccharide transporters (Doidy et al., 2012), just a few reacted on the presence of mycorrhizal colonisation – namely the probes for one tonoplast monosaccharide transporter (TMT) and two poyol/monosaccharide transporters (PMT), named here PMTa and PMTb (see Table S1 for further details).

**TMT** was significantly downregulated at two time points in M+ roots (Fig. S4). Unlike most of the other genes of interest, which were almost not-expressed in shoots, TMT showed relatively high level of expression in the shoots, where it was upregulated at 35 dpp but downregulated at 49 and 63 dpp. However, in Exp 2, the downregulation of TMT was not significant. Upon 90% shading, M+ roots displayed significantly lower expression of TMT after 4 days of shading, although no change was detected in the NM roots (Fig. S4). Regarding shaded shoots, TMT expression was not significantly altered in the M+ shoots. In contrast, 90% shaded NM shoots upregulated the expression of TMT at both time points.

A “switch-like” character of expression was shown for **PMTa**, which was upregulated at 35 and 49 dpp in M+ roots, but suddenly and strongly increasing the expression in the NM roots (Fig. S4). Interestingly, the expression differences were not confirmed in Exp 2 (Fig. S4). Also, the upregulation of PMTa in shaded roots (both M+ and NM) was not strongly supported by the statistics. In shoots, PMTa was only little expressed (tens of copies per 1 µg of RNA; Fig. S4).

Activation of **PMTb** in M+ roots was clearly visible (Fig. S4), however, its upregulation at 49 dpp was not significant due to big dispersion of data. Nevertheless, Exp 2 confirmed its upregulation in M+ roots (Fig. 3). Furthemore, PMTb was downregulated in M+ roots upon shading, with a drop to NM levels after 4 days of shading. In both M+ and NM shoots, the expression of PMTb was on comparable level as is in the roots. Moreover, it was upregulated once in the M+ shoots (63 dpp), and downregulated once upon 90% shading in the M+ shoots.

**Figure 3:**
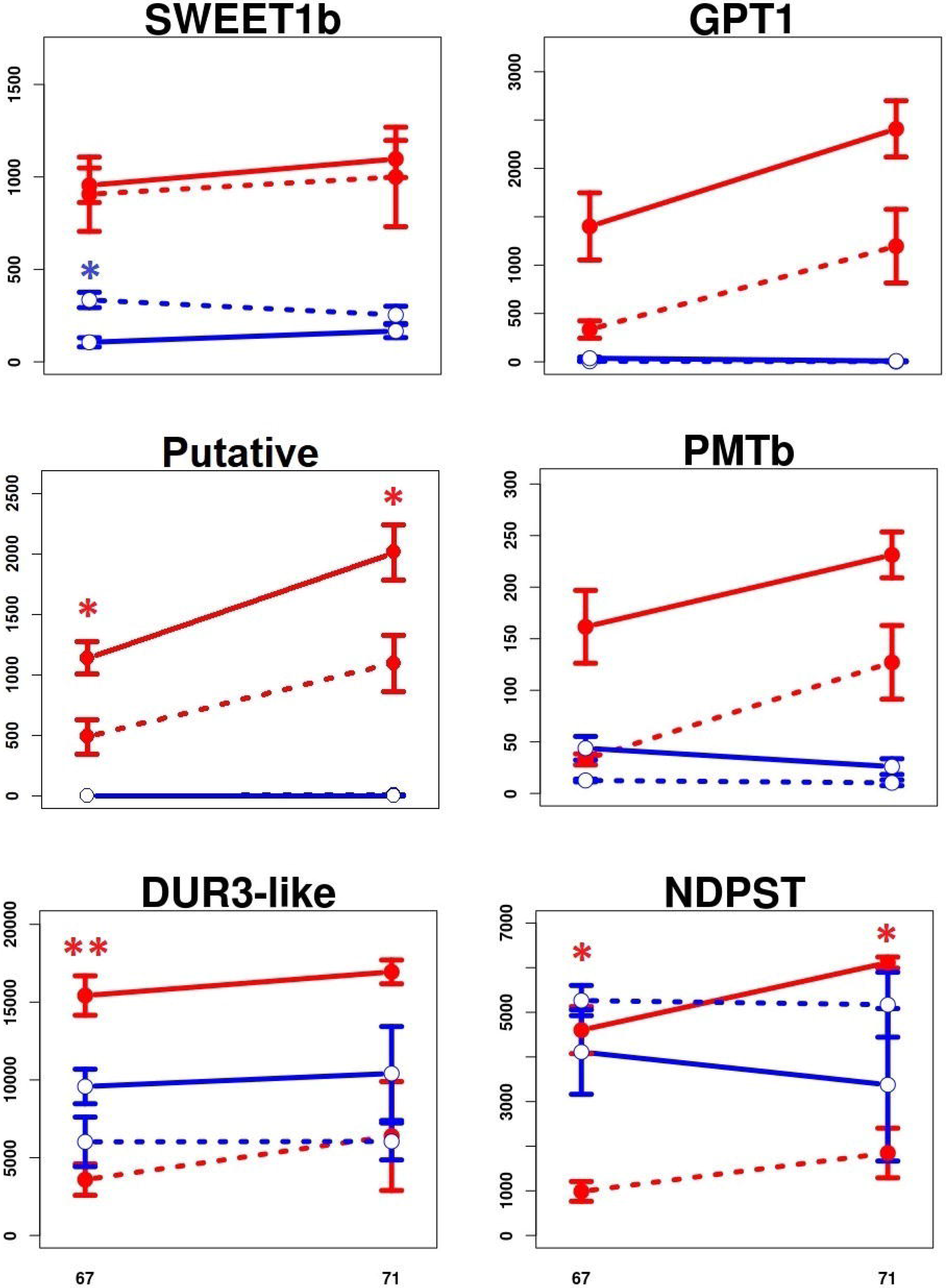
The expression of several genes obtained by qPCR from roots of M. truncatula in Exp 2. X-axis: days post planting, Y-axis: number of corresponding-gene copies per 1 µg of RNA subjected to reverse transcription. Red: mycorrhizal, Blue: non-mycorrhizal, Full line: full light (100%), dashed-line: shaded plants (10% of light). Error bars show standard deviation, n = 3. For further details see Materials and methods and Table S1. Signif. codes for t-test between non-/shaded: 0 < *** < 0.001 ≤ ** < 0.01 ≤ * < 0.05 ≤ non-significant.

In another search of keyword “sugar” using application “Samples” (Genevestigator), 113 probes with matching anotation were found, and another 3 genes encoding saccharide transpoters with altered AM-induced expression were selected: two bidirectional sugar transporters SWEET1b and SWEET3a, and a sugar porter Mtst1.

While the upregulation of **SWEET3a** in M+ roots was evident based on the microarray data (Table S1), significant difference between the M+ and NM roots was not confirmed by qPCR (Fig. S4). Moreover, at 63 dpp, SWEET3a was expressed more in the NM roots. Upon 90% shading in Exp 2, the expression in the M+ roots decreased, but only after 4 days of shading the results significantly differed between full light and shaded treatments (Fig. S4).

The expression of **SWEET1b** appeared to be modulated by the AMS, too – its expression in NM roots was low and the upregulation in M+ roots was statisticaly significant in all time points included in Exp 1 (Fig. S4). Yet, the expression of the gene seems not to have reacted to shading of the plants in M+ roots, whereas in NM roots the shading upregulated its expression transiently (Fig. 3). In shoots, the expression levels of this gene were close to zero (Fig. S4).

The expression of **Mtst1** (Harrison, 1996) displayed a similar “switch-like” pattern as the PMTa did – it was upregulated in M+ roots at 35 dpp, then the differences disappeared and finally, at 63 dpp, Mtst1 was downregulated (Fig. S4). In Exp 2, the expression of Mtst1 was more or less similar. In shoots, the gene was only very slighty expressed (tens of copies per 1 µg of RNA; Fig. S4).

With a similar approach, using keyword “glucose”, a strong shift in expression level of glucose-6-phosphate / phosphate translocator 1 (**GPT**) gene located on chromosome 8 was found. While almost not-expressed in NM plants, we saw a clear upregulation of GPT in M+ roots (Fig. S4) and in reponse to short-term shading, the expression levels decreased significantly (Fig. 3). Its expression in shoots was close to zero (Fig. S4).

### DUR3-like, NDPST and putative transporter – novel genes in arbuscular mycorrhizal symbiosis

Also some other genes somewhat connected to saccharide metabolism or transportome of the plants were selected, based on keywords “symport”, “sugar” or “transmembrane”, and their differential expression in the M+ and NM plants. Their expression level in M+ roots promised a relevance to the AMS, with direct or indirect relevance to the transfer of C from the plant to the AM fungus.

Strong upregulation of **DUR3-like** encoding gene at 35 dpp in M+ roots pointed towards a specific role of this transporter in M+ roots, though it probably symports Na^+^ and urea. The difference in expression of this gene between M+ and NM roots disappeared at 49 dpp, but it was upregulated again at 63, 67 and 71 dpp (Fig. S4 and Fig. 3). In shoots, DUR3-like encoding gene was not expressed very strongly. Yet, upon shading, its expression levels dropped in both M+ and NM shoots, indicating its potential involvement in shoot metabolism no matter the symbiosis (Fig. S4).

The only enzyme (not transporter) examined in this study was a nucleotide-diphosphosugar transferase (**NDPST**). Its expression seemed to be upregulated in M+ roots, but it dropped at 49 dpp (Fig. S4) and the pattern was not the same in Exp 1 and Exp 2 (Fig. 3) – however, upon shading, the expression decrease was conspicious. There was no significant difference between M+ and NM plants in the NDPST expression in shoots (Fig. S4).

Last but not least, one putative membrane transporter was selected in this study (named **Putative** here). Similarly to GPT and PMTb, it was almost not expressed in NM roots and both M+ and NM shoots (Fig. S4), but its expression was strongly upregulated in M+ roots (Fig. S4). More importantly, upon shading, it was significantly downregulated in M+ roots (Fig. 3). Based on topology prediction, using HMMTOP, CCTOP and PredictProtein, 12 transmembrane helices were predicted (data not shown), which is a common feature of ST of major facilitator superfamily (Yan, 2013). It is predicted to localize into endoplasmic reticulum, but the PredictProtein do not count with PAM. Most importantly, BLASTp search in non-host species’ database showed no results (see Discusion for further information).

### The expression of GPT, PMTb, DUR3-like, NDPST and Putative transporter correlates with expression of PT4 in M+ roots upon shading

In Exp 2, the expression levels of PT4 decreased significantly in response to the shading of the plants. We used this induced response to correlate expressional changes of other genes tested in this study in M+ roots with those of PT4. Several genes displayed very similar pattern of downregulation upon shading as the PT4 gene (Fig. 4), while the rest of examined genes showed no correlation (data not shown). For shoot samples, no correlations were evaluated.

**Figure 4:**
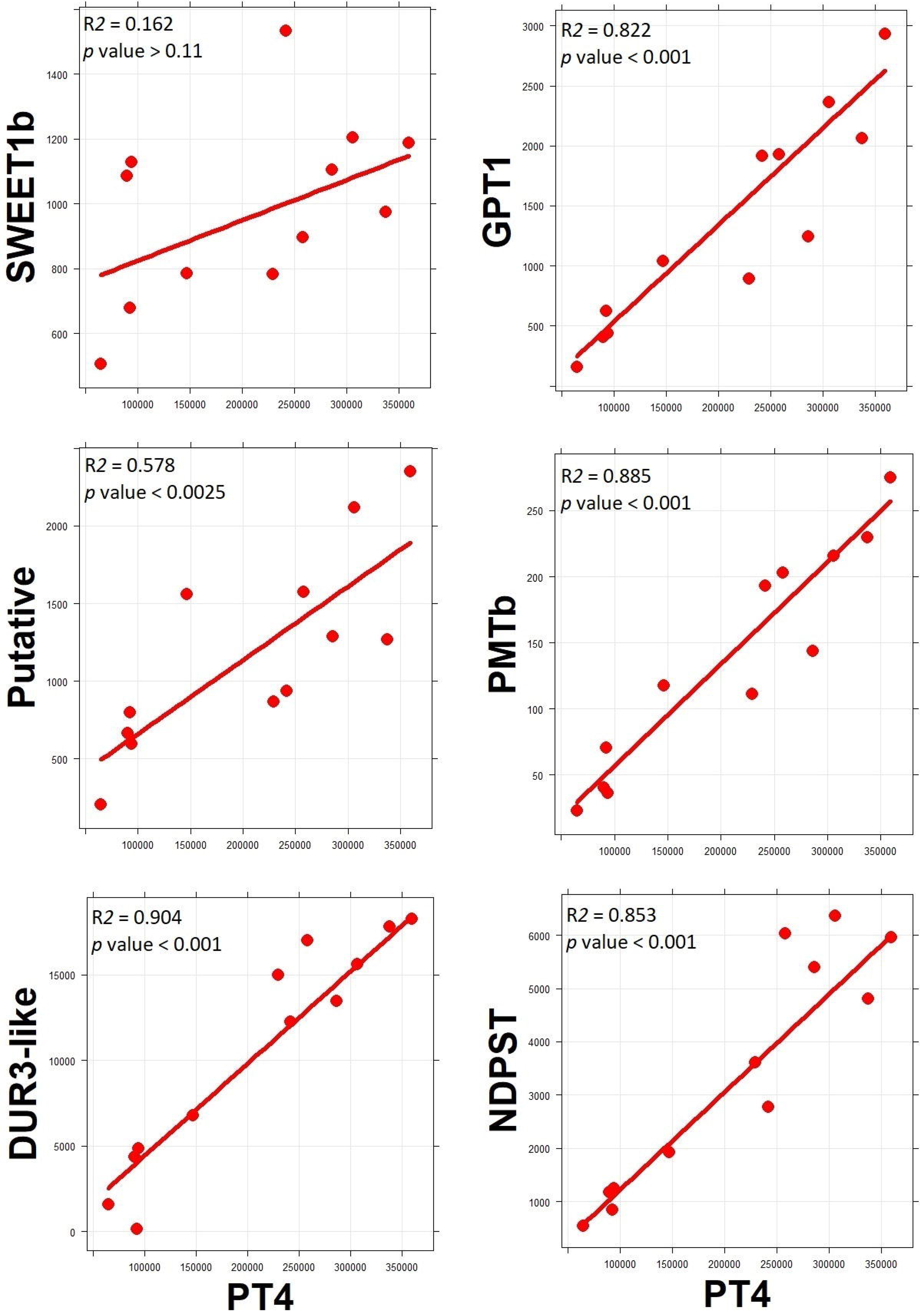
Scatterplots with simple linear regression of expressions of several genes against PT4 expression of mycorrhizal root samples in Exp 2. X-axis: number of PT4 copies per 1 µg of RNA subjected to reverse transcription, Y-axis: number of corresponding-genes copies per 1 µg of RNA subjected to reverse transcription. Both harvests (67 and 71 dpp) and both light treatments (100% and 10% light) are taken as a single group.

The SWEET1b gene surprisingly did not change its expression in response to shading, which was further supported by *p* value greater than 0.11 for simple linear correlation with PT expression in the M+ roots. In contrary, the simple linear correlations of GPT, PMTb, DUR3-like, NDPST and Putative transporter expressions with the expression of PT4 were all significant (Fig. 4). Since the differences in expression of those genes in roots of M+ shaded and non-shaded plants were mostly significant (Fig. 3), we assume a direct involvement of these genes in AMS, at least regarding their transcriptional regulation.

In terms of the expression patterns of the above genes, there are three distinct patterns (Fig. 3). For GPT1 and Putative transporter genes the situation was identical as for PT4: in NM plants, the genes were virtually not expressed, but in M+ plants they were expressed and downregulated by shading (Fig. 4). For PMTb and DUR3-like genes, the situation with respect to response to shading in M+ plants was analogous to GPT1 and Putative: the genes were significantly downregulated upon shading in M+ plants and the correlation with PT4 was significant, too. However, in contrast to GPT1 and Putative transporter, PMTb and DUR3-like genes were also expressed in the NM plants, yet in a generally lower level than in the M+ plants. Finally, NDPST gene was expressed similarly in both NM and M+ plants exposed to full light, but upon shading, only in M+ roots it was (significantly) downregulated (Fig. 3) and well correlated with PT4 expression (Fig. 4).

## Discussion

In this study we grew barrel medic (*Medicago truncatula*) with compatible rhizobia and with or without AM fungus *Rhizophagus intraradices*. This common AMS model system was used in two separate experiments, Exp 1 and Exp 2, to identify novel genes possibly involved in shuffling reduced C ompounds from the plant to the AM fungus (Fig. 1). AMS was fully established in the M+ plants, as shown by AM colonization (Fig. S3), and by the expression of AM-specific PT4 gene (Fig. 2). Oppositely, the NM plants remained free from the AM fungal colonization. Interestingly, the plants of both treatments were similar in terms of biomass production (Fig. S1), whereas the M+ plants had twice as high P concentration as the NM plants (Fig. S2). This indicates a functional mycorrhizal P uptake pathway, and (also because of lack of the positive growth response of the plants to AMS formation), also an extensive trading of P for C between the symbionts (Lendenmann et al., 2011, Řezáčová et al., 2018), presumably at the periarbuscular interface. For deciphering a transcriptional regulation of the pre-selected genes possibly invoved in the C transfer from the plant to the AM fungus, we grew the plants for roughly a couple of weeks at full light, and then applied a short-term shading treatment as suggested previously (Konvalinková et al., 2015).

We have identified a transcriptional changes in two SWEET genes of *Medicago truncatula* – SWEET1b and SWEET3a. SWEET proteins are pH-independent, bidirectional ST with broad representation in bacteria, plants, fungi and animals (Chen et al., 2010; Chen et al., 2012; Lin et al., 2014) with crucial roles in multiple biological processes. **SWEET1b** shows upregulation in M+ plants, similar to the rice gene OsSWEET1b (Samueelah et al., 2016) or potato genes StSWEET1a and StSWEET1b (Manck-Götzenberger & Requena, 2016). Closest characterized homologue is AtSWEET1 of thale cress (*Arabidopsis thaliana*), whose substrate specificity is higher for Glc and somewhat lower for Gal, but does not transfer Suc (Chen et al., 2010), so similar substrate specificity could be expected for **SWEET1b** as well. Usually, the homologues are expressed in sink tissues like fruit, nodules, tubers or upon mycorrhization, so the role in apoplastic transport of Glc is assumed. Notably, if **SWEET1b** localizes on PAM, it allows Glc to leak out, where RiMST of the *R. irregularis* takes up the Glc molecules immediately, and simultaneously, it serves as passive retractor of Glc to the plant cells, with following utilisation of Glc in plant cytoplasm or sequestring into a vacuole (Fig. 5). The fungal ST posses high-affinity to Glc – RiMST2 has a K_m_ of 33 ± 12.5 μM (Helber et al., 2011), and AtSWEET1 is a low-affinity glucose transporter with a K_m_ of 9 mM (Chen et al., 2010). That means, the net efflux of Glc from arbuscocyte is expected. On the other hand, the Glc retracted from periarbuscular space by SWEET1b will be actively utilised by cytoplasmic hexokinase to Glc-6-P, and since the hexokinase has a K_m_ of 20-130 μM for Glc (Claeyssen and Rivoal, 2007), the competition for Glc between plant and fungus may occur even when no active ST is located at the PAM (Fig. 5).

**Figure 5:**
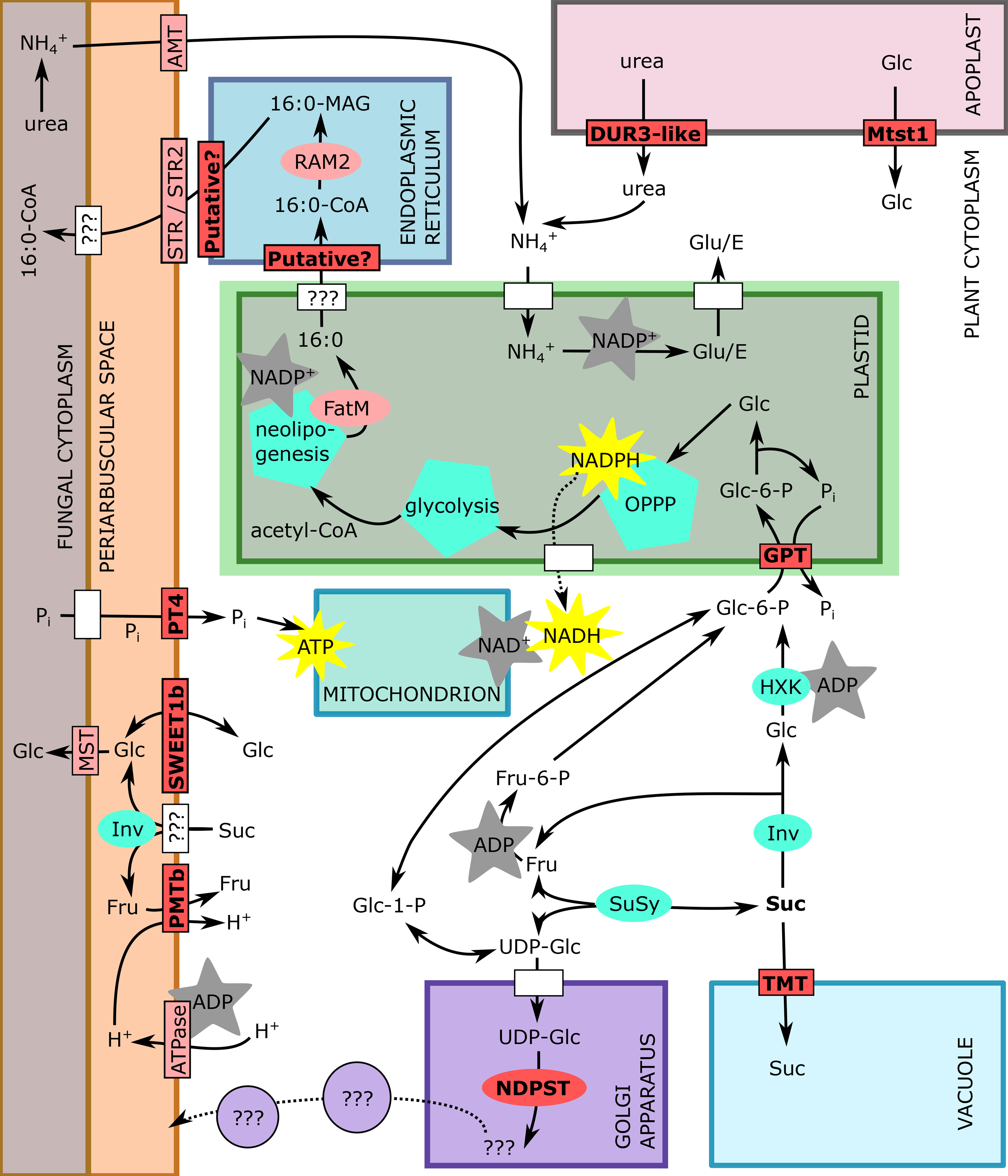
Scheme of arbuscocyte showing investigated genes (bright red), and some of AM-relevant proteins (light red), enzymes and enzymatic pathways (cyan), and not specified proteins (white). Generation of energy molecules (ATP) and reductive coenzymes (NADH / NADPH) in yellow star, while consuming of those in gray star. 16:0, free palmitoyl; 16:0-CoA, palmitoyl-Coenzyme A; 16:0-MAG, mono acyl glycerol (palmitoyl glycerol); acetyl-CoA, acetyl Coenzyme A; UDP-Glc, uridine diphosphate glucose; Glc-1-P, glucose-1-phosphate; Glc-6-P, glucose-6-phosphate; Fru-6-P, fructose-6-phosphate; P_i_, phosphate; Inv, invertase; SuSy, sucrose synthase; Suc, sucrose; Glc, glucose; Fru, fructose; HXK, hexokinase; OPPP, oxidative pentose phosphate pathway; Glu/E, glutamate;

Another AM-specific candidate gene is **GPT**. This gene has longer mRNA (contrary to the other 2 GPT genes in *Medicago truncatula* genome) and is annotated also as O-glycosyl hydrolase. Since no substrate specificity has been establisled for this gene with respect to hydrolysis, this enzymatic annotation needs to be further investigated. GPT in general imports Glc-6-P to plastids in exchange of P to maintain starch synthesis, oxidative pentose-phosphate pathway (OPPP), neolipogenesis or nitrite reduction, mainly in heterotrophic tissues (Kammerer et al., 1998). This crossroad of cell metabolism secures C flow to plastid, while enabling sequestration towards storage (anabolism) or katabolism and symbiont-feeding by lipids (Fig. 5). In plastids, products of OPPP may serve for shikimate pathway producing aromates, for methylerithritolphosphate pathway (MEPP) producing isoprenoids, for synthesis of nucleotides, and other molecules like phytohormones (Lohse et al., 2005). Assuming that all these metabolic pathways are already active in NM cells, more or less, this GPT can hardly be the main C importer of plastids, since we measured almost zero expression in the NM plant roots (Fig. 3 and Fig. S4). On the other hand, we show that GPT is an AM-specific gene, which is regulated in the same manner as PT4. It would be interesting to further investigate if GPT expression and / or activity is directly linked to presumed lipid-transfer machinery like STR/STR2 (Gutjahr et al., 2012; Rich et al., 2017b).

**Putative** membrane transporter (MTR_8g071050) was chosen because it is predicted to posses 12 transmembrane helices (data not shown) and this is a common feature of proteins from Major Facilitator Superfamily (Yan, 2013). We present three arguments for involvement of this Putative membrane transporter in AMS. First, we find almost zero expression in NM roots, so it is an AM-induced gene (Fig. 3 and Fig. S4). Second, its downregulation in shaded M+ roots is significantly correlated with the downregulation of PT4, so it is AM-controled gene (Fig. 4). Third, simple BLASTp search (https://blast.ncbi.nlm.nih.gov/Blast.cgi) of its protein sequence (XP_013446273.1) results in hits in broad range of land plant families, but no hit can be found when narrowing the query to *Amaranthaceae* or *Brassicaceae* family only (data not shown), which are known NM families (Harley and Harley, 1987). Recently, it has been shown that phylogenomic approach is useful for identification of AMS-conserved genes, when the orthologues are present in AM-species and absent in NM-species (Bravo et al., 2016). Therefore, we assume that this uncharacterized Putative membrane transporter is recruited specifically for AMS action.

When we look on **PMTb** gene probe in the gene expression atlas (MtGEA; https://mtgea.noble.org/v3/), it is expressed in nodules, and there is no expression pattern recorded of **PMTb** in the AMS. But nodules were present in all treatments (although we did not quantify the nodule formation in our experiments, so quantitative differences may have escaped our attention). Closest *A*. *t*. homologues symports Fru and xylitol with H^+^ so it activelly imports Fru from apoplast after cleavage of Suc (Klepek et al., 2010). Small change in **PMTb** activity on PAM should be sufficient for actively resorb Fru, because even a small amount of transporters may effectively change the concentrations of solutes in the periarbuscular space (Schott et al., 2016). However, since **PMTb** is not expressed in AM roots without nodules, the precise localization and function of this gene is remaining an unsolved puzzle.

The homologue of **DUR3-like** gene in *Arabidopsis thaliana*, AtDUR3, is a symporter of urea and sodium ions (Liu et al., 2003) and it was described to mediate the retrieval of urea from senescensing *Arabidopsis leaves‘* apoplast (Bohner et al., 2015). The periarbuscular space is a type of apoplastic compartment, so the urea could be taken up by DUR3-like, but since the exchange of NH_4_^+^ between the plant and the fungus is fairly described (Jin et al., 2012; Calabrese et al., 2016; Wang et al., 2017), this option is unlikely. More likely, the urea derives from senescent structures like root cells or degenerated arbuscules, since the arbuscules are ephemeral structures (Luginbuehl and Oldroyd, 2017).

**NDPST** glycosylates its substrates and this one NDPST (out of 23 described NDPST *M*. *t*. genes) is annotated as glucosylating by UDP-Glc. Does it posses a role in glucosylation of membrane lipids or proteins? The best possibility is an involvement in cell wall component synthesis in Golgi apparatus (Fig. 5) – the cell wall biogenesis (Sakiroglu and Brummer, 2016). But since there is no degradation of plant cell wall polymers by AM fungi (Tisserant et al., 2013; Balestrini & Bonfante, 2014), there is probably no reason for delivering cell wall polymers into periarbuscular space. In this study, we have discovered a strong evidence for AM-dependent regulation of NDPST, which displays a similar pattern as the regulation of PT4 – responding a short-term shading of the plants. Another NDPST (MTR_8g069400) is predicted to be AM-conserved gene (Bravo et al., 2016), so the relevance of (a change of) glycosylation upon AMS is fair. Interestingly, the lipid pathway was depicted as a variation of cutin biosynthesis pathway (Rich et al., 2017a), and higher production of cell wall components is a common defence strategy against fungal pathogens (Underwood, 2012; Bellincampi et al., 2014).

Paszkowski et al. (2002) writes: “Thus, although cytological and physiological features of the arbuscular mycorrhizal symbiosis seem to be conserved, the molecular components may differ significantly between distantly related plant species. “ Since the properties of ST, like substrate specificity, transport activity or localization may differ much between different plants, while sequence comparisons show sometimes strong similarities, the next step for deciphering sugar partitioning in AM plants is the biochemical characterization and subcellular localization of plant ST during AMS. There definitely is still much to be done with respect to full undestanding of the molecular mechanisms of bidirectional trading of resources in the AMS, particularly with respect of C flux from plants to the AM fungus, including different C forms (Keymar and Gutjahr, 2018). Yet this appears necessary to achieve the next break-through in understanding of functioning of this ancient interkingdom relationship (Parniske, 2008) and possibly its better utilization for human welfare (Bender et al., 2015; van de Wiel 2016).

## Abbreviations

qPCR: quantitative polymerase chain reaction
PT: phosphate transporter
ST: saccharide transporter
GPT: glucose-6-phosphate/phosphate translocator
G6P: glucose-6-phosphate
Mt: Medicago truncatula
AM: arbuscular mycorrhizal
C: carbon
N: nitrogen
P: phosphorus
M+: mycorrhizal treatment
NM: non-mycorrhizal treatment
RPM: rotations per minute
DEPC: diethylpyrocarbonate
SDS: sodiumdodecylsulphate
EDTA: ethylenediaminetetraacetic acid
EtOH: ethanol
RNase: ribonuclease
DNase: deoxyribonuclease
RNA: ribonucleic acid
DNA: deoxyribonucleic acid
cDNA: complementary DNA
Suc: sucrose
Glc: glucose
Fru: fructose
PAM: periarbuscular membrane
dpp: days post planting
RT: reverse transcription
AMS: arbuscular mycorrhizal symbiosis

## Author Contributions

Both JK and JJ planned the experiments and interpreted the results, JK, HH, PB and MH conducted the laboratory analysis, JK carried out *in silico* part of the work, then both JK and JJ jointly wrote the manuscript and all authors agreed with the manuscript in its final form.

## Acknowledgments

The authors wish to thank Hana Gryndlerová, Tereza Konvalinková, Václav Šťovíček, Veronika Procházková and Milan Gryndler from Laboratory of Fungal Biology for watering the plants and kind help with harvests, and Osaro Konečná for graphical assistance.

## Data Availability Statement

The experimental data are deposited in NCBI’s Gene Expression Omnibus (Edgar et al., 2002) and are accessible through GEO Series accession number GSE126833 (https://www.ncbi.nlm.nih.gov/geo/query/acc.cgi?acc=GSE126833) as mentioned in Materials and methods.

