## Supplemental Figures for "Correlative evidence for co-regulation of phosphorus and carbon exchanges with symbiotic fungus in the arbuscular mycorrhizal *Medicago truncatula*"

### Figure S1

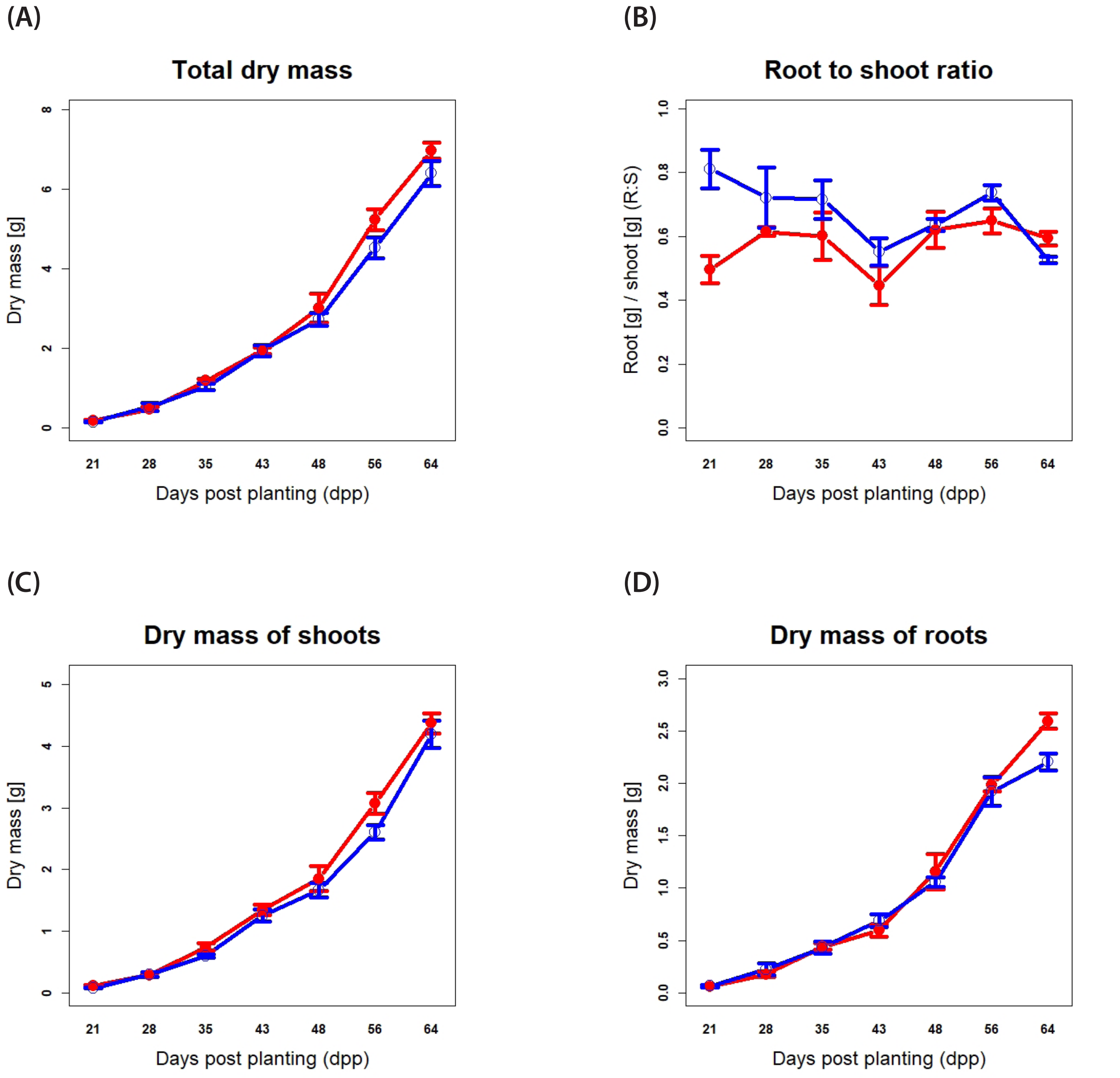


**Figure S1:** Dry mass of M. truncatula in Exp 1. **(A)** total dry mass of whole plants; **(B)** root to shoot ratio; **(C)** dry mass of whole shoots; **(D)** dry mass of whole roots; Error bars show standard deviation, n = 3. **Red:** mycorrhizal, **Blue:** non-mycorrhizal. Signif. codes for t-test between M+/NM: 0 < *** < 0.001 ≤ ** < 0.01 ≤ * < 0.05 ≤ non-significant.

### Figure S2

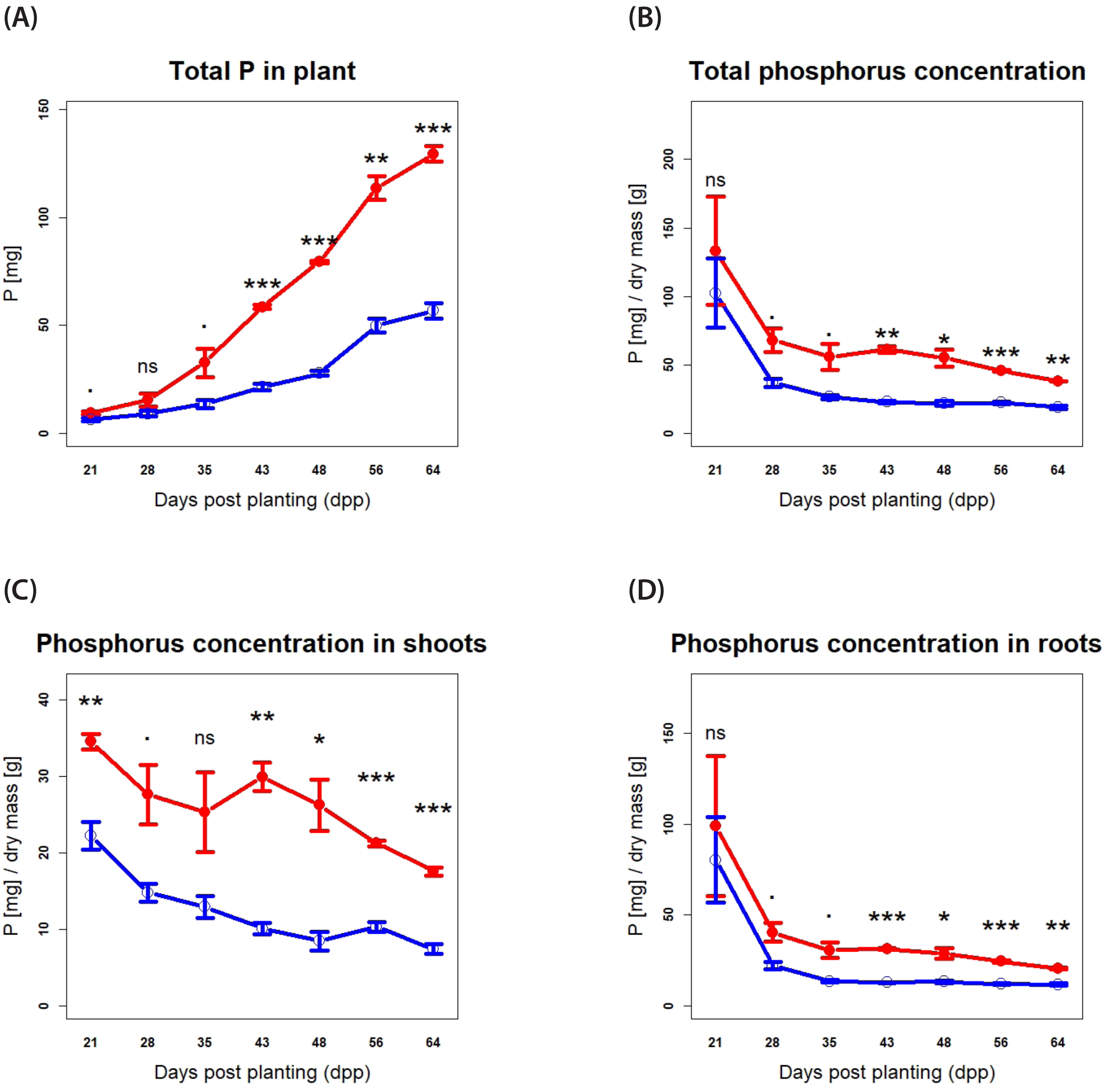


**Figure S2:** Evaluation of phosphorus (P) content in M. truncatula in Exp 1. **(A)** Total P content in whole plants; **(B)** total P concentration in whole plants; **(C)** P concentration in whole shoots; **(D)** P concentration in whole roots; Error bars show standard deviation, n = 3. **Red:** mycorrhizal, **Blue:** non-mycorrhizal. Signif. codes for t-test between M+/NM: 0 < *** < 0.001 ≤ ** < 0.01 ≤ * < 0.05 ≤ non-significant.

### Figure S3

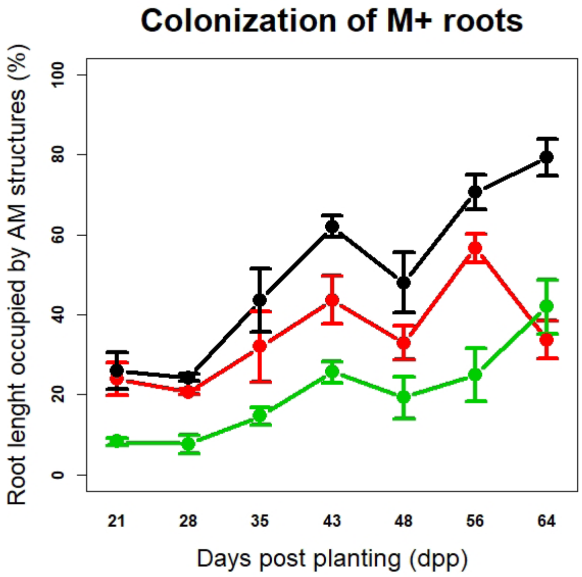


**Figure S3:** Evaluation of mycorrhizal colonization of mycorrhizal roots of M. truncatula in Exp 1. Error bars show standard deviation, n = 3. **Black:** hyphae; **Red:** arbuscules; **Green:** vesicles.

### Figure S4

**(A) Exp 1 - roots**

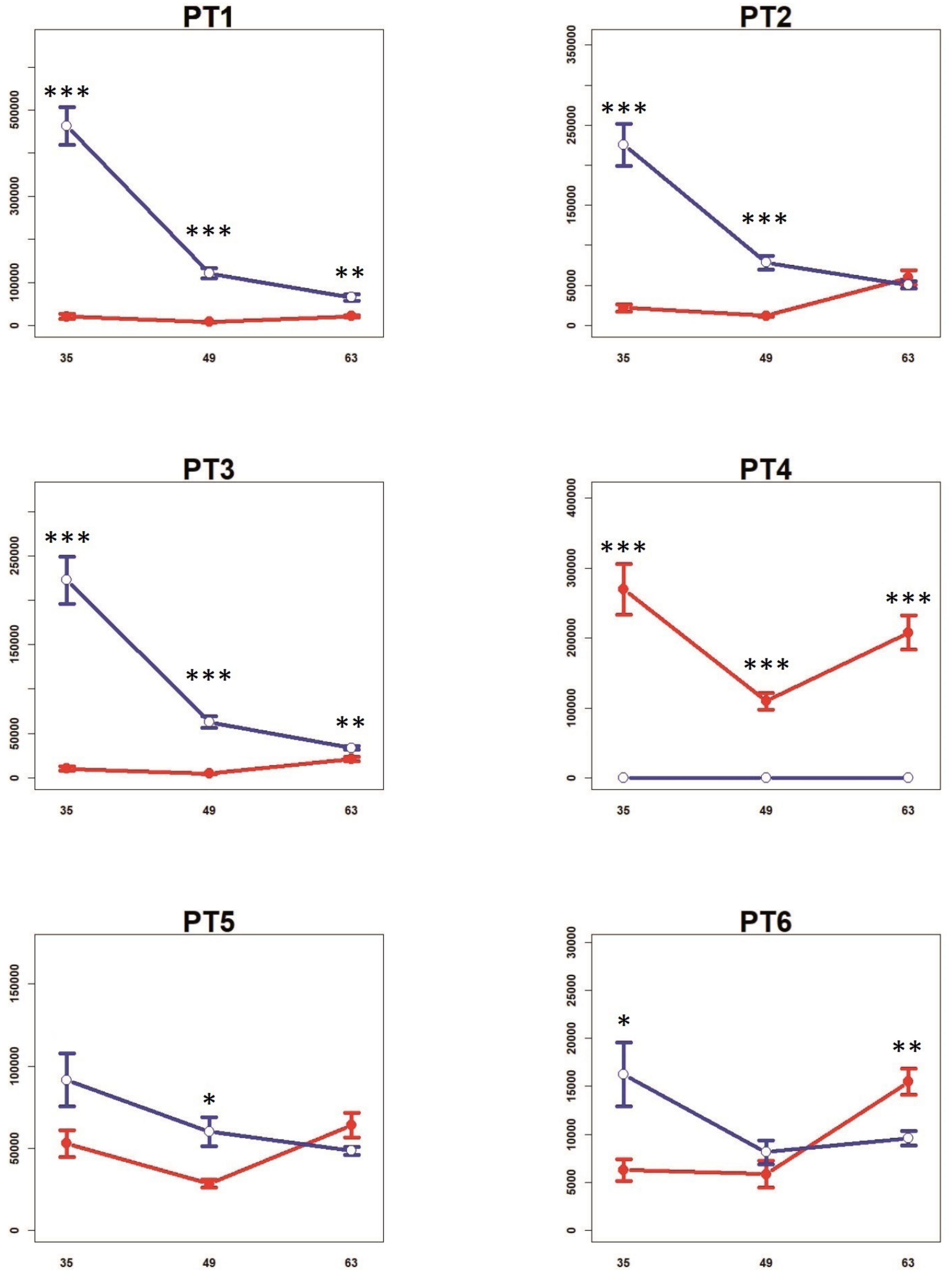

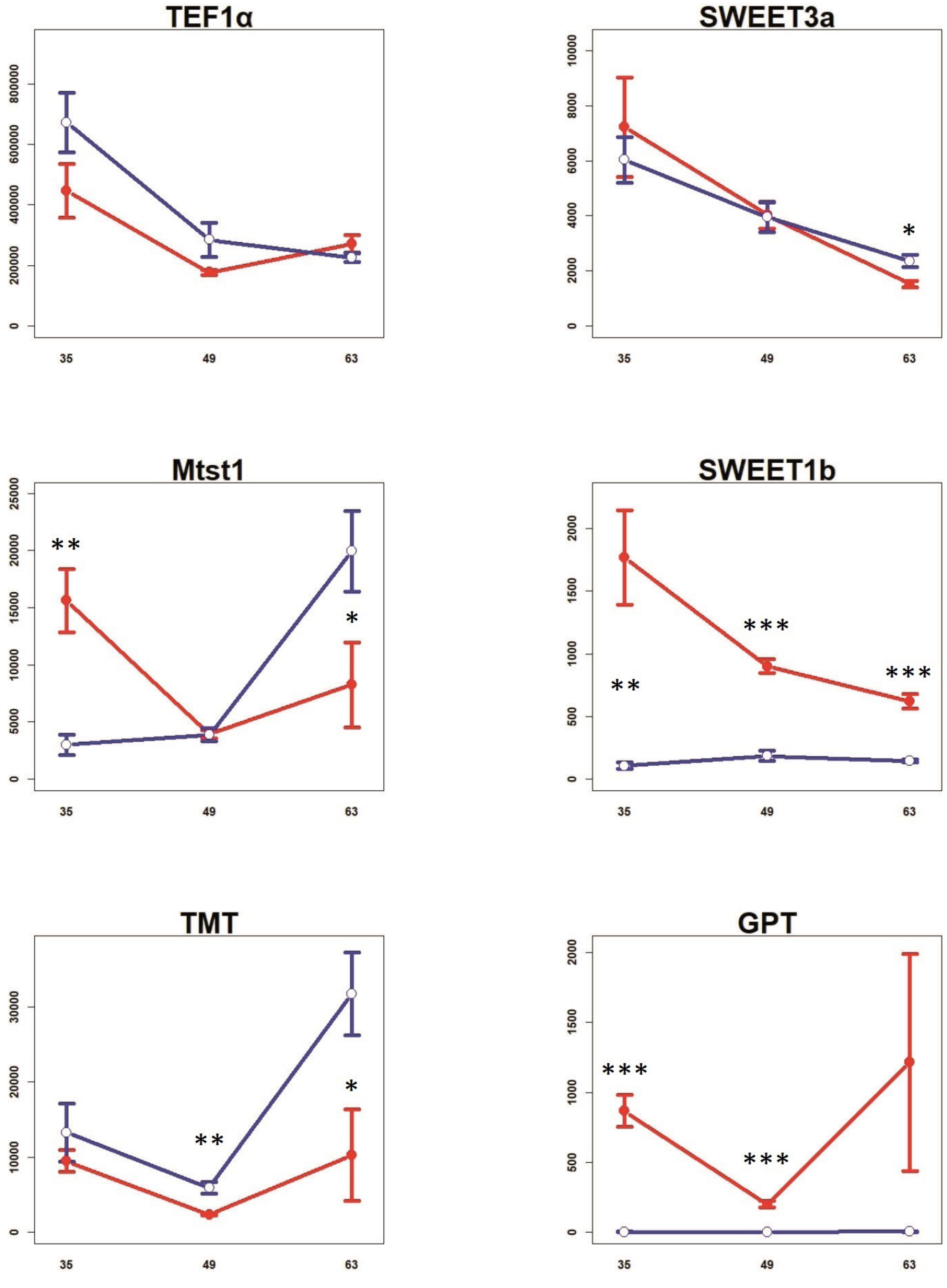
TEF1α
0
200000
400000
600000
800000
35
49
63
SWEET3a
0
2000
4000
6000
8000
10000
*
35
49
63
Mtst1
0
5000
10000
15000
20000
25000
**
*
35
49
63
SWEET1b
0
500
1000
1500
2000
**
***
***
35
49
63
TMT
0
10000
20000
30000
**
*
35
49
63
GPT
0
500
1000
1500
2000
***
***
35
49
63

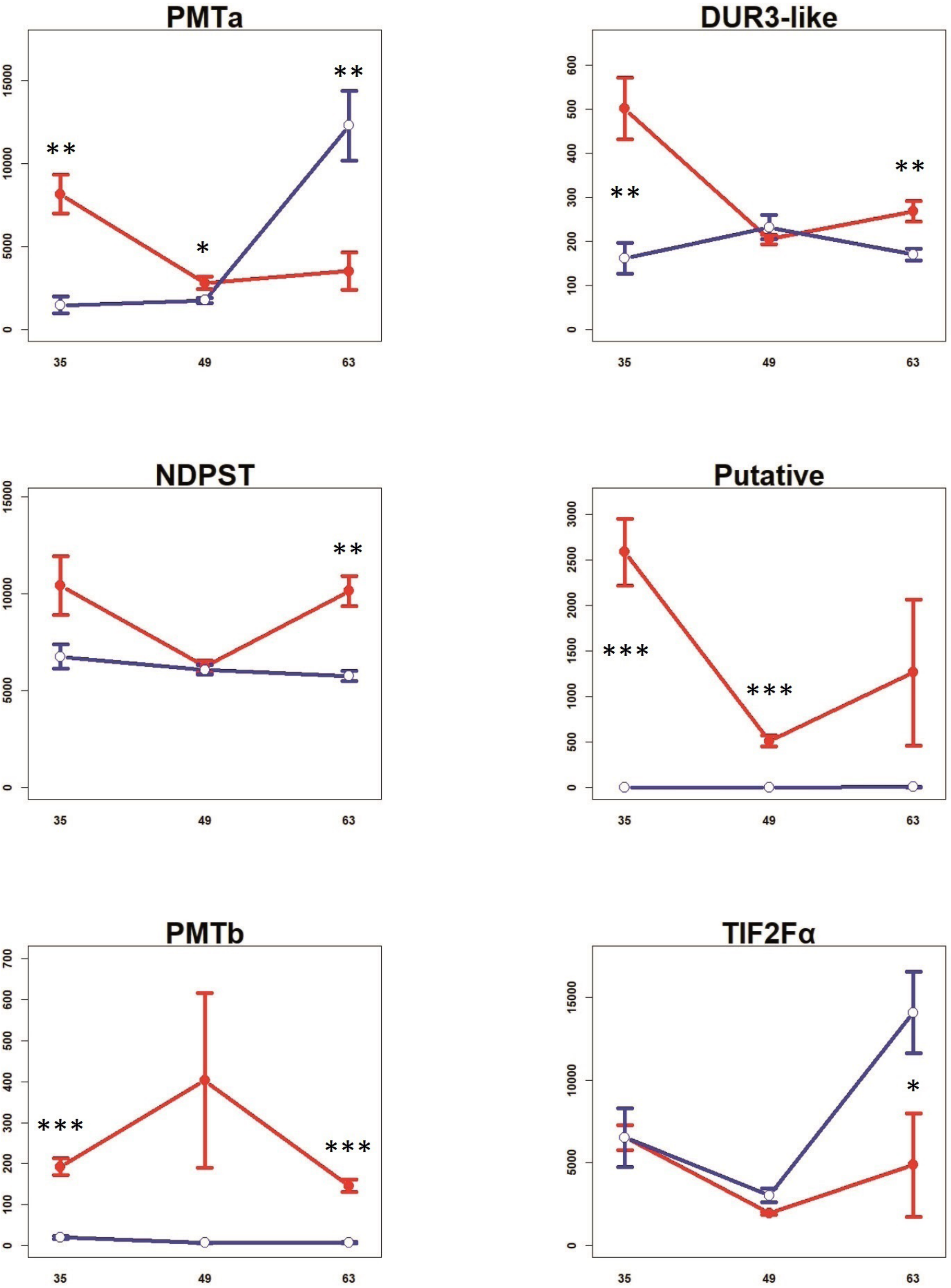

PMTa
DUR3-like
NDPST
Putative
PMTb
TIF2Fα
35
49
63

**(B) Exp 1 - shoots**

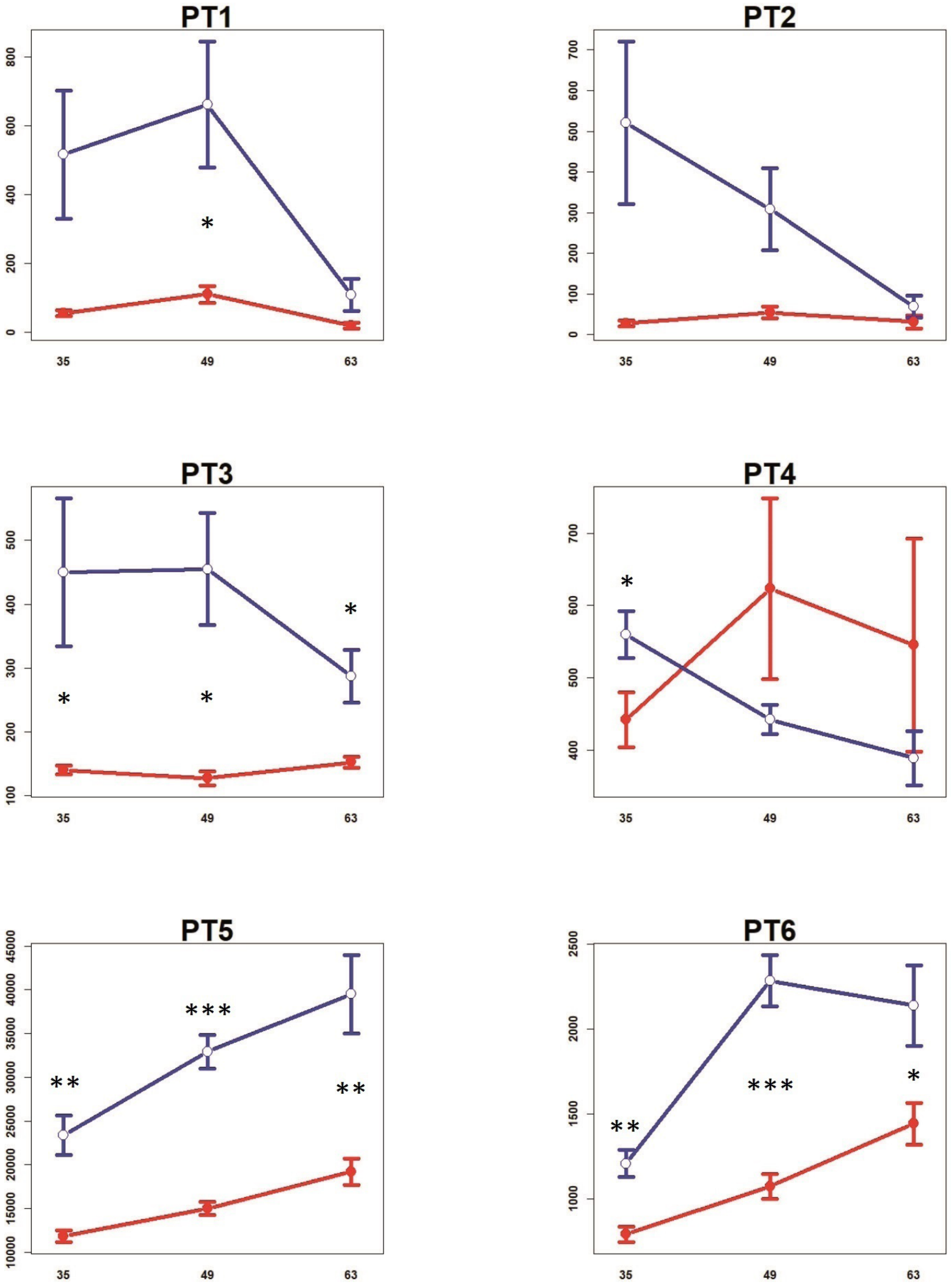

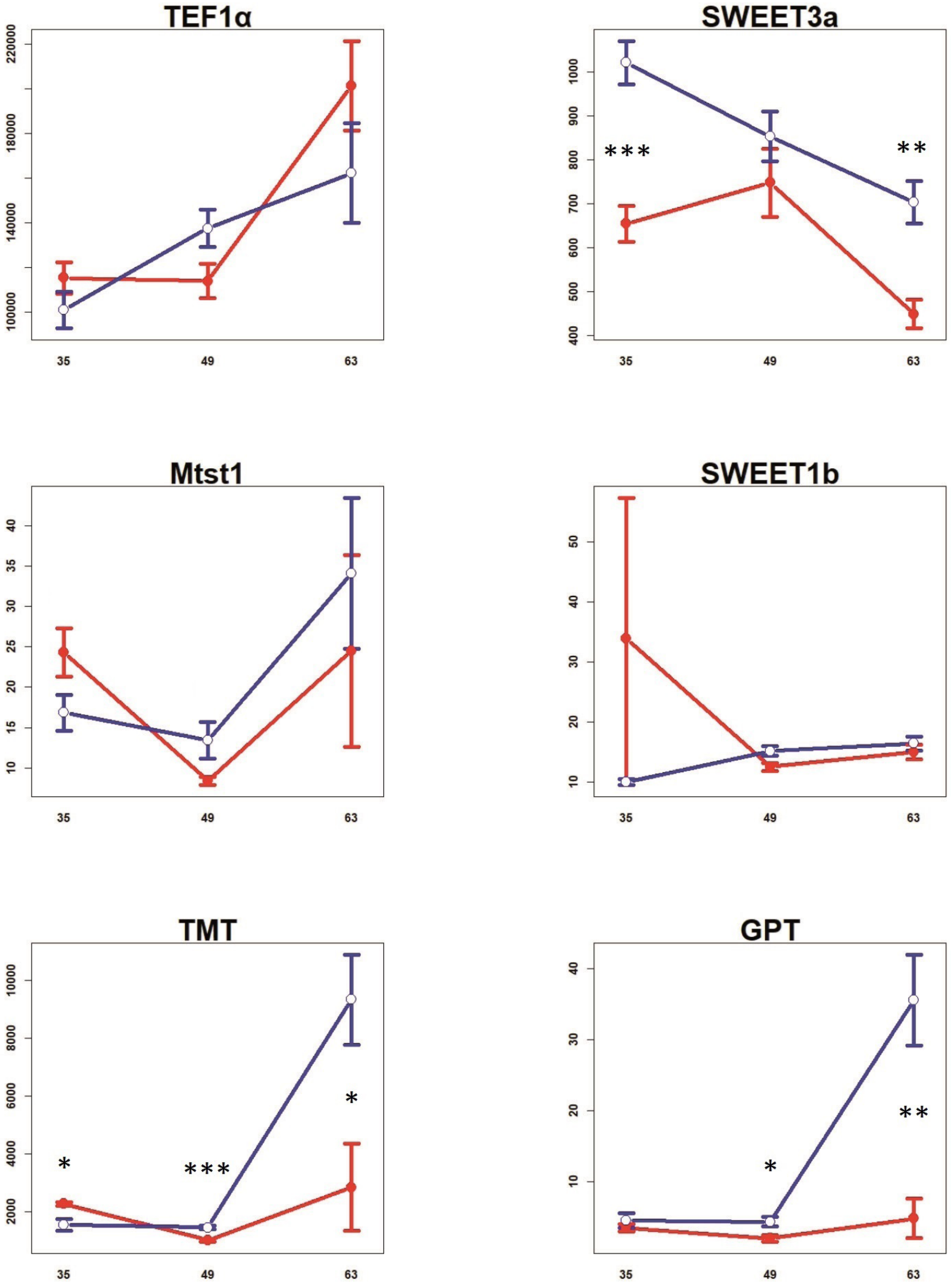
TEF1α
220000
180000
140000
100000
35
49
63
SWEET3a
1000
900
800
700
600
500
400
***
**
35
49
63
Mtst1
40
35
30
25
20
15
10
35
49
63
SWEET1b
50
40
30
20
10
35
49
63
TMT
10000
8000
6000
4000
2000
*
***
*
35
49
63
GPT
40
30
20
10
*
**
35
49
63

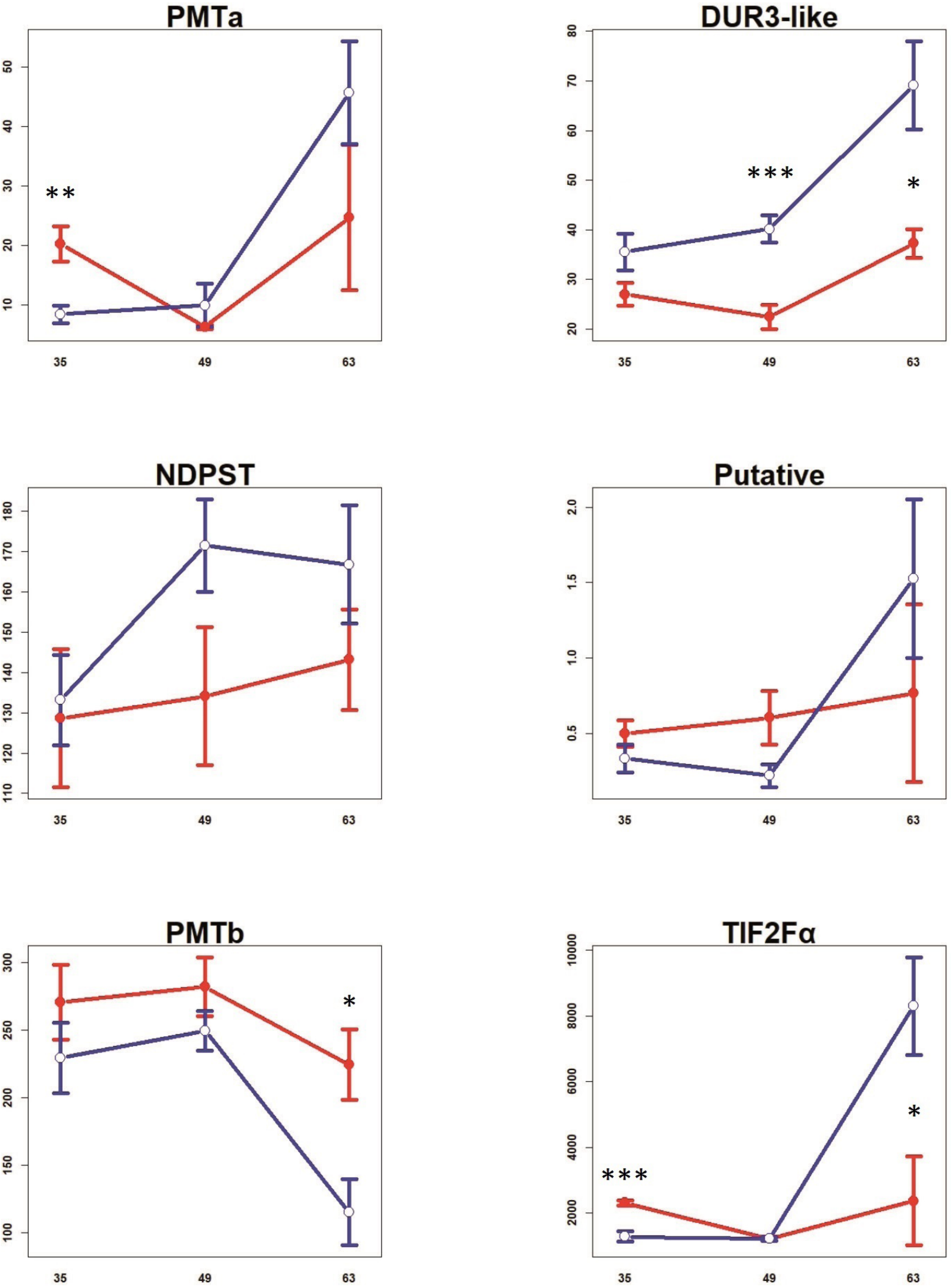
PMTa
DUR3-like
NDPST
Putative
PMTb
TIF2Fα
**
***
*
***
*
*
35
49
63

(C) Exp 2 - roots

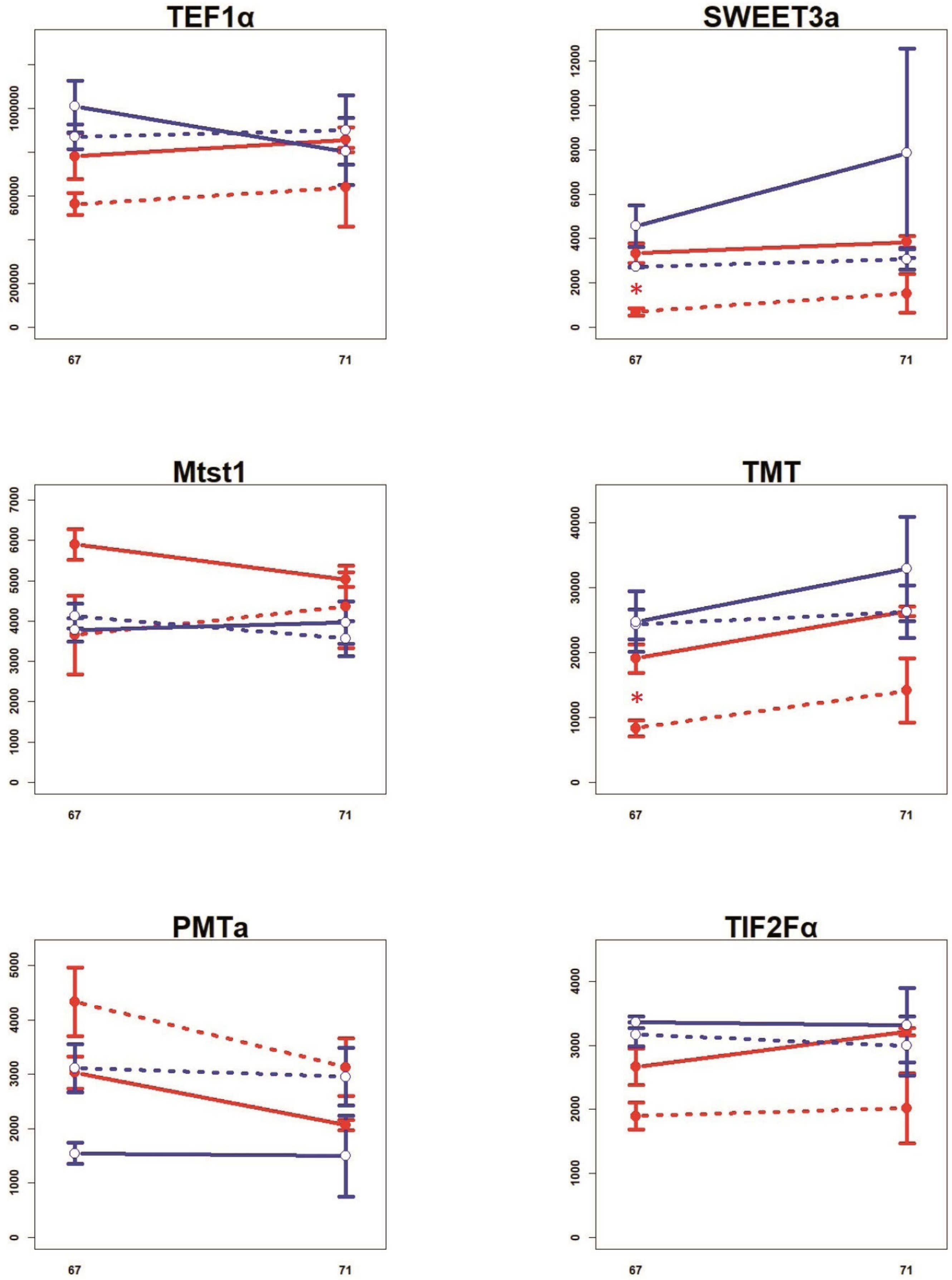
TEF1α
0
200000
600000
1000000
67
71
SWEET3a
0
2000
4000
6000
8000
10000
12000
*
67
71
Mtst1
0
1000
2000
3000
4000
5000
6000
7000
67
71
TMT
0
10000
20000
30000
40000
*
67
71
PMTa
0
1000
2000
3000
4000
5000
67
71
TIF2Fα
0
1000
2000
3000
4000
67
71

(D) Exp 2 - shoots

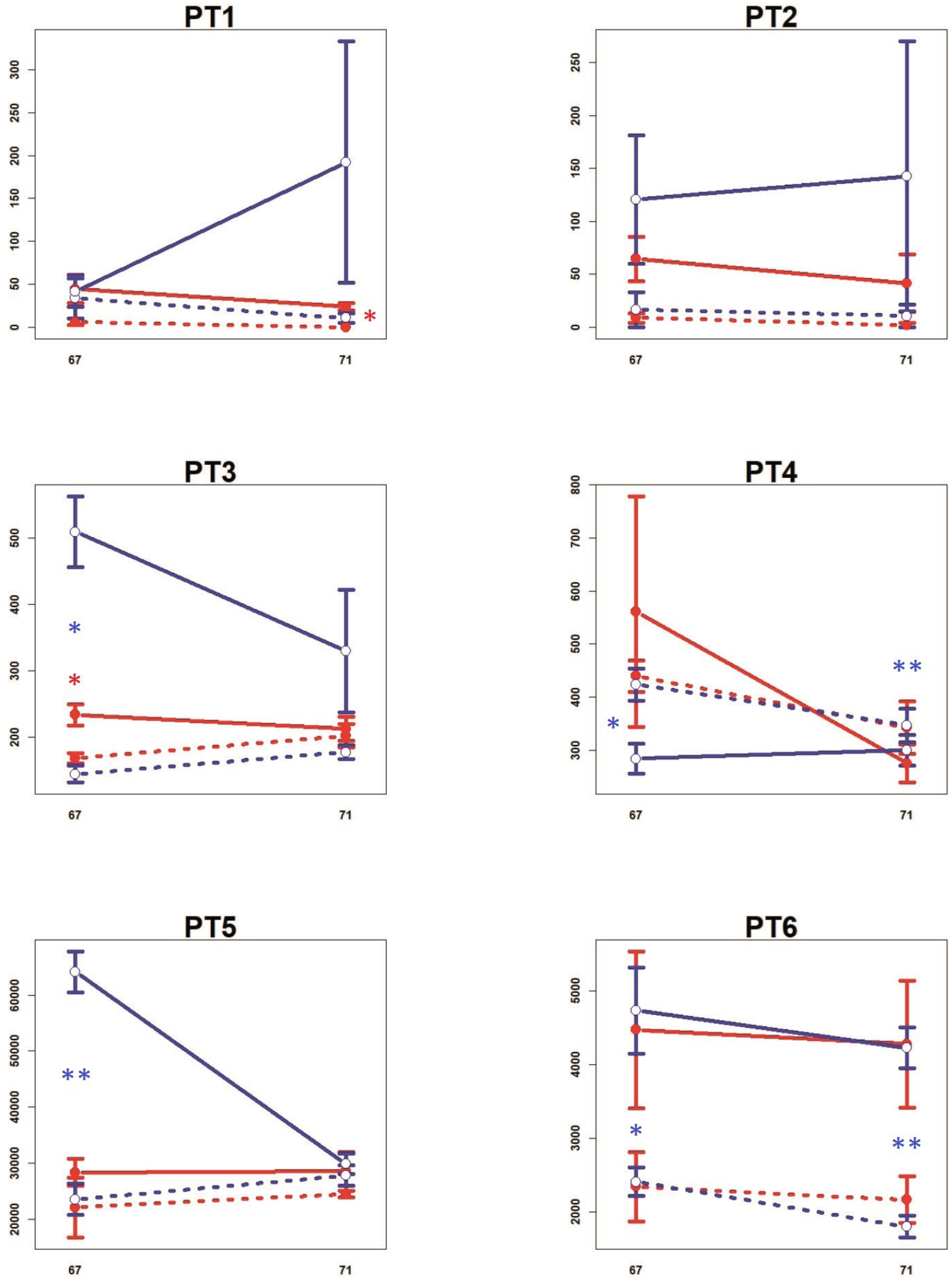

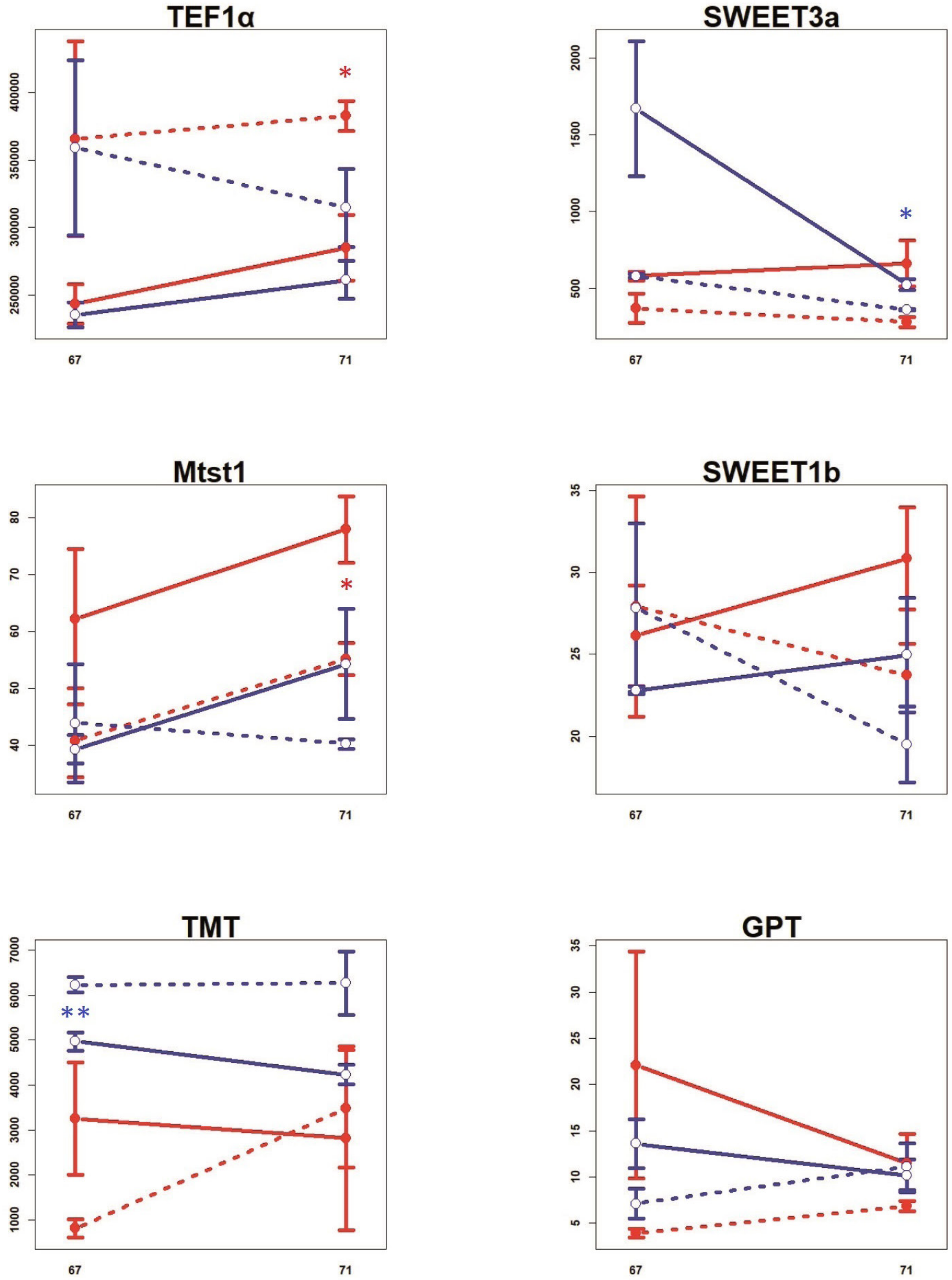
TEF1α
400000
350000
300000
250000
*
67
71
SWEET3a
2000
1500
1000
500
*
67
71
Mtst1
80
70
60
50
40
*
67
71
SWEET1b
35
30
25
20
67
71
TMT
7000
6000
5000
4000
3000
2000
1000
**
67
71
GPT
35
30
25
20
15
10
5
67
71

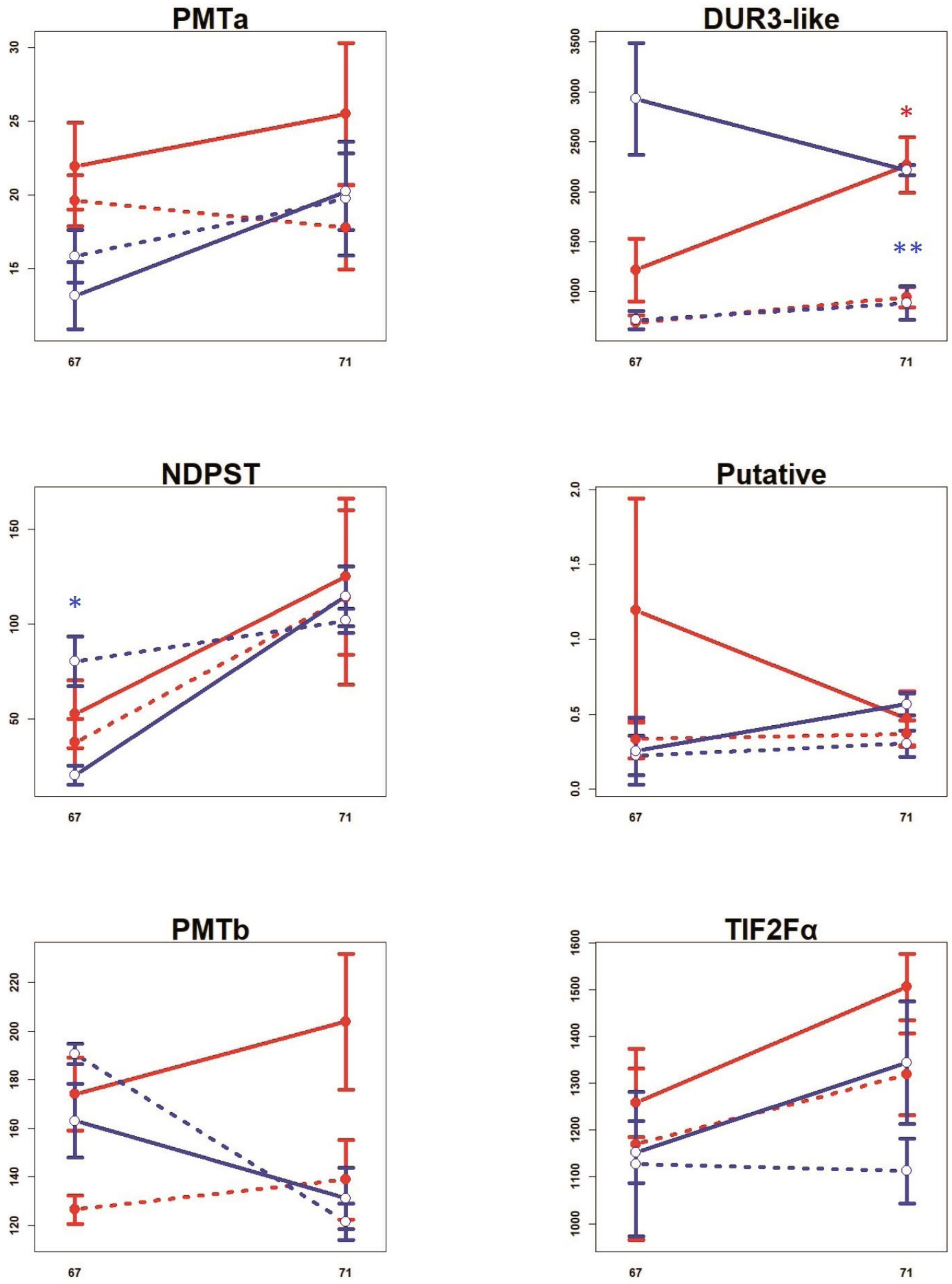


**Figure S4:** The expression of genes obtained by qPCR from of M. truncatula roots **(A and C)** or shoots **(B and D)** in Exp 1 **(A and B)** or Exp 2 **(C and D)**, respectively. X-axis: days post planting, Y-axis: number of corresponding-gene copies per 1 microg of RNA subjected to reverse transcription. **Red:** mycorrhizal, **Blue:** non-mycorrhizal; Full line: full light (100%), dashed-line: shaded plants (10% of light). Error bars show standard deviation, n = 6 or 3 for Exp 1 or Exp 2, respectively. For further details see Materials and methods and Table S1. Signif. codes for t-test between full light/shaded plants: 0 < *** < 0.001 ≤ ** < 0.01 ≤ * < 0.05 ≤ non-significant. For "missing" graphs see Fig. 2 and Fig. 3.
